# Cooperative IDR-folded domain interplay governs HP1α phase separation and underlies biphasic regulation by pathogenic poly-PR

**DOI:** 10.64898/2026.10.10.758178

**Authors:** Yize Zhang, Dan Su, Huanyu Li, Hongfeng Wang, Juan Li, Chaowei Shi, Tie Shen, Zhonghuai Hou, Zheng Ying, ShengQi Xiang

## Abstract

HP1α undergoes liquid-liquid phase separation (LLPS), yet the prevailing model, which attributes condensation almost entirely to intrinsically disordered regions (IDRs), leaves the role of its folded domains unresolved. Using solution and solid-state NMR, we show that HP1α LLPS is instead governed by reciprocal interplay between IDRs and folded domains. Within the condensed phase, solid-state NMR reveals a dynamic hierarchy: the chromoshadow domain (CSD) assembles into polymorphic oligomers that form rigid cores, while the phosphorylation sites on the N-terminal extension (NTE) remain highly mobile, challenging their proposed role as stable multivalent hubs. Phosphorylation of the NTE instead promotes LLPS primarily by increasing local rigidity and releasing the NTE from the chromodomain (CD) to relieve autoinhibition, with electrostatic charge contributing secondarily. The C-terminal extension (CTE) suppresses phase separation through two routes: weakening the association between the hinge region (HR) and the CSD, and inhibiting interactions between CSD dimers. ALS-associated poly-PR targets this network at the NTE and exerts concentration-dependent biphasic regulation, promoting LLPS at low concentrations via multivalent bridging and dissolving condensates at high concentrations via saturation, a switch recapitulated in cells and in mouse brain. These findings reframe HP1α condensation from an IDR-driven process to one cooperatively governed by IDR-folded domain interplay.

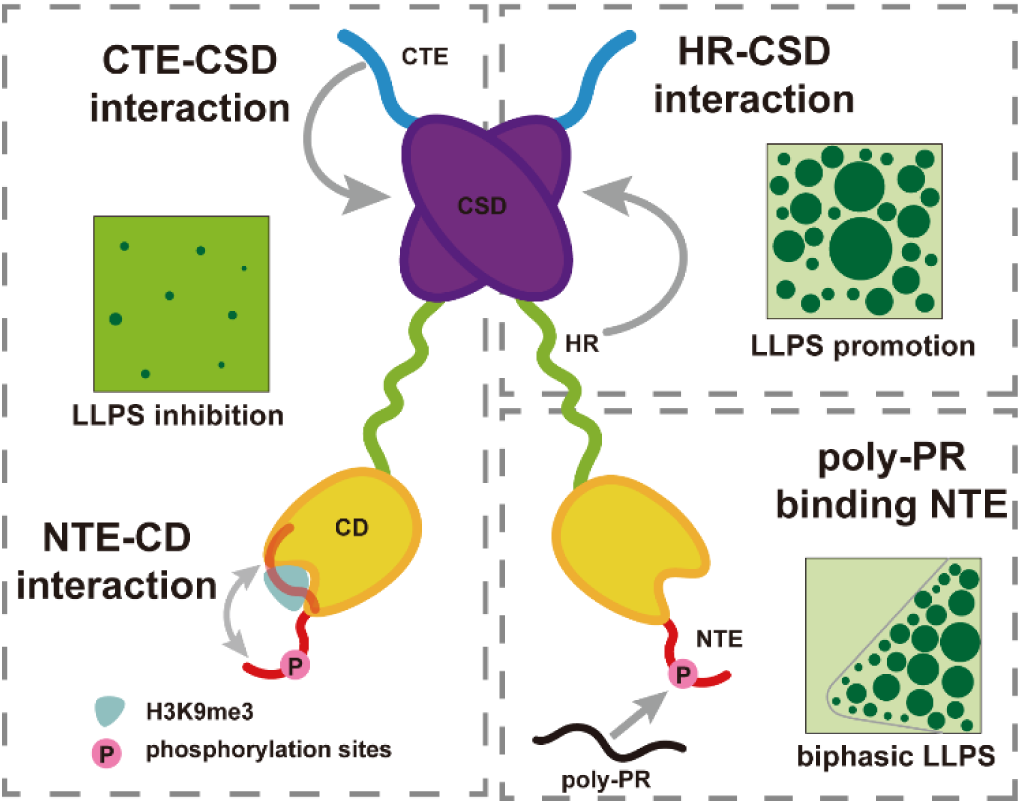

## Introduction

HP1α is a central effector of heterochromatin, driving the assembly and maintenance of repressive chromatin domains [1-4]. This function rests on a multi-domain architecture (Fig. 1A) consisting of three intrinsically disordered regions (IDRs): the N-terminal extension (NTE), hinge region (HR), and C-terminal extension (CTE), and two highly conserved folded domains: the chromodomain (CD) and the chromoshadow domain (CSD). The CD recognizes H3K9me3, a repressive histone mark [5-8], while the CSD mediates homodimerization and PXVXL-motif ligand recognition for protein recruitment [9-11]; the CSD also harbors additional interfaces enabling higher-order oligomerization, promoted by exogenous KAP1Hbox [12-14]. The NTE is phosphorylated at four consecutive serines, thereby increasing the negative charge and extending the conformation of the HP1α dimer [15-17]. The positively charged HR serves as a central hub for interactions, engaging the NTE, CTE, and nucleic acids [16, 18-22]. MD simulations further suggest that the CTE may modulate the CSD oligomerization interface [13]. Notably, the intramolecular contacts defined to date converge on the disordered HR, while the folded domains are known mainly as ligand-binding modules; whether they also engage in the intramolecular interaction network remains unexplored.

**Fig. 1.**
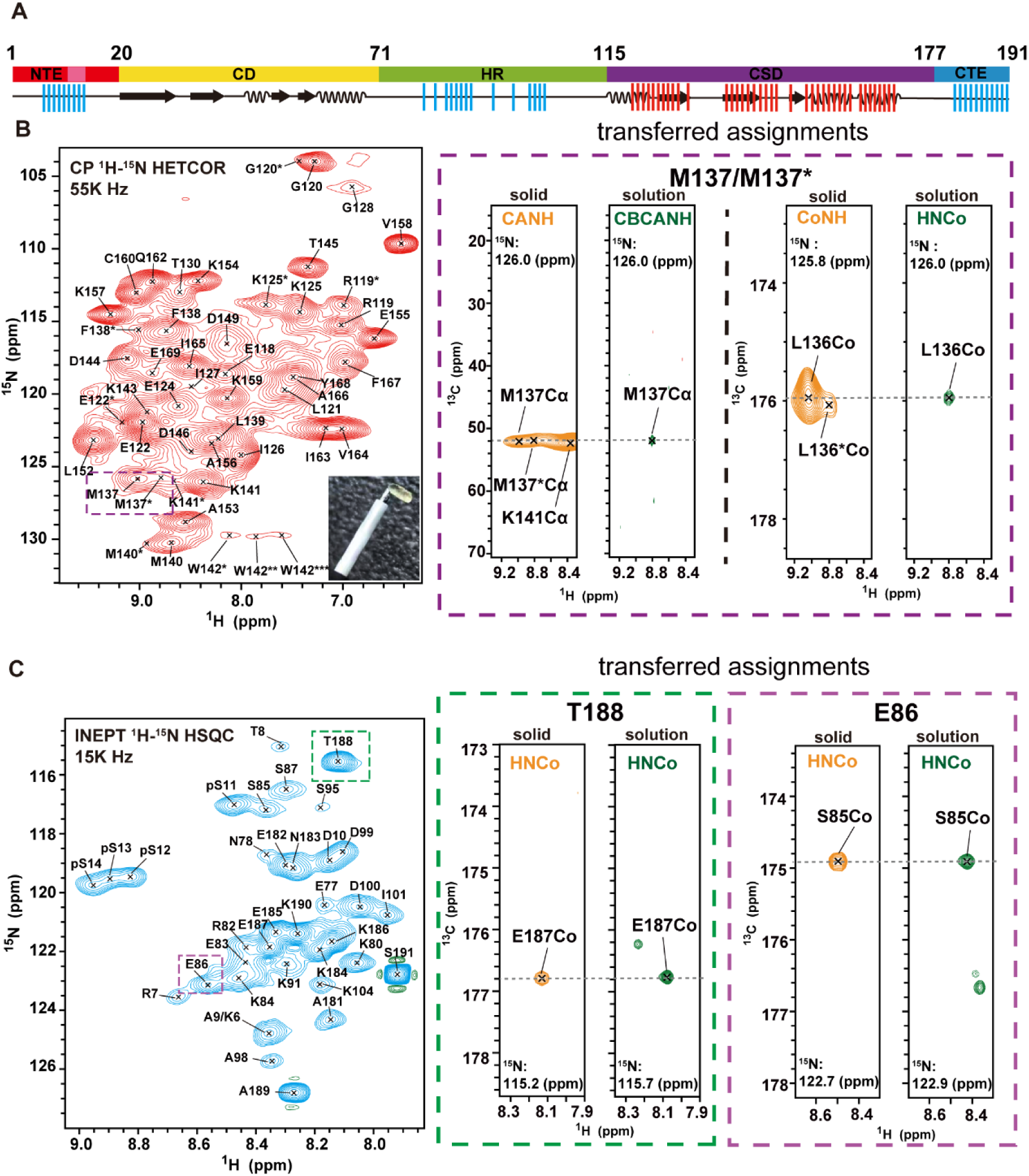
Solid-state NMR reveals distinct domain dynamics within pHP1α condensates. **(A)** Domain architecture of HP1α. Secondary structure elements are shown below the linear domain map. Residues detected in the CP-based spectrum (Fig. 1B) are highlighted in red, and residues detected in the INEPT-based spectrum (Fig. 1C) are highlighted in blue. NTE, N-terminal extension; CD, chromodomain; HR, hinge region; CSD, chromoshadow domain; CTE, C-terminal extension. **(B)** CP-based ¹H-¹⁵N HETCOR solid-state NMR of condensed pHP1α detects rigid domains. Strong signals are observed exclusively from the CSD; CD signals are absent. Insets show transferred assignments and multiple chemical shifts for M137, indicating polymorphic CSD oligomers. The 1.3 mm rotor and the condensed phase are shown at the bottom right of the ¹H-¹⁵N HETCOR spectrum. **(C)** INEPT ¹H-¹⁵N HSQC solid-state NMR detects flexible IDRs within pHP1α condensates. Phosphorylated serines yield well-resolved signals, indicating high local dynamics. Insets show transferred assignments for representative residues.

HP1α undergoes liquid-liquid phase separation (LLPS) both in vivo and in vitro [16, 23-25]. The prevailing model posits a simple electrostatic logic: NTE phosphorylation increases negative charge, thereby enhancing multivalent interactions with the positively charged HR to form an IDR-driven network that drives LLPS [16, 17, 19, 26, 27]; conversely, the CTE is thought to suppress phase separation by competing with the NTE for HR binding [16, 20, 28, 29]. This framework treats the folded CD and CSD largely as passive scaffolds, responsible for histone recognition and dimerization, respectively, while assigning the driving force of condensation almost entirely to the IDRs.

However, this charge-centric, IDR-dominant model suffers from several major inconsistencies. First, phosphomimetic NTE mutants (e.g., SSSS→EEEE) fail to promote phase separation in vitro [16], and charge-neutralizing NTE mutations do not prevent HP1α foci formation in vivo [17], indicating that changing charge alone is insufficient to regulate condensation. Second, CTE deletion strongly enhances phase separation even in the absence of NTE phosphorylation [16, 18], suggesting that CTE suppression is not solely mediated by competition with NTE for HR binding but likely involves additional mechanisms that remain undefined. Third, how CD-bound H3K9me3 promotes phase separation remains unclear within the prevailing framework [16]. Moreover, although MD simulations suggest that folded domains may also contribute to phase separation, the precise mechanisms remain to be experimentally verified [20]. Meanwhile, previous solid-state NMR studies of HP1α condensates lacked residue-specific assignments [30-33], precluding atomic-resolution analysis of the condensed phase, and computational predictions of contact patterns within the condensed phase lack direct experimental validation [20, 28].

HP1α condensation is also a target of disease-associated perturbation. ALS-associated poly-PR dipeptide repeats, produced by C9orf72 repeat-associated non-AUG translation [34-37], disrupt HP1α LLPS compartments and compromise heterochromatin integrity[38]. Poly-PR is a highly cationic, multivalent repeat polymer known to broadly perturb biomolecular condensates [39], yet the molecular basis of its action on HP1α remains unresolved.

Here, using solution and solid-state NMR, we dissect the IDR-folded domain interaction network governing HP1α phase separation at residue resolution. Within condensates, solid-state NMR reveals a dynamic hierarchy: the CSD assembles into polymorphic oligomers that form rigid cores, whereas the phosphorylation sites on the NTE remain highly mobile, challenging their proposed role as stable multivalent hubs. We find that NTE phosphorylation promotes phase separation primarily by increasing local rigidity and releasing the NTE from the CD, thereby relieving autoinhibition; charge plays only a secondary role. We experimentally demonstrate that CTE suppresses phase separation not only by blocking the HR-CSD interaction, but also by directly binding the CSD to inhibit dimer-dimer interactions. ALS-associated poly-PR binds NTE and exerts biphasic regulation, promoting LLPS at low concentrations via multivalent bridging and dissolving condensates at high concentrations via saturation, with both regimes recapitulated in cultured cells. These findings reframe HP1α condensation from an IDR-centric process to one governed by cooperative interplay between IDRs and folded domains.

## Results

### Solid-state NMR reveals structural and dynamic heterogeneity within pHP1α condensates

Despite extensive studies on HP1α LLPS, the internal structural organization and residue-level dynamic behavior of its folded domains and IDRs within the condensate remain poorly characterized. We therefore induced pHP1α LLPS to form stable condensates and performed the residue-level solid-state NMR analysis of the condensed phase. The pHP1α sample was verified by solution-state ¹H-¹⁵N HSQC to carry complete phosphorylation at all four NTE serines (S11-S14; Fig. S1), ensuring a homogeneous phosphorylation state for the analyses below. Using cross-polarization (CP)-based ¹H-¹⁵N HETCOR [40-43], we detected signals from rigid, motion-restricted structured domains (Fig. 1B); INEPT ¹H-¹⁵N HSQC captured signals from flexible IDRs (Fig. 1C) [42, 44]. We further acquired three-dimensional CP and INEPT spectra to obtain Cα and Co chemical shifts in the condensed phase, transferring assignments from solution-state NMR (Fig. 1A-C).

With these assignments, we observed a striking internal dynamic hierarchy within the condensate. In the INEPT ¹H-¹⁵N HSQC experiment, the four phosphorylated serines yielded well-resolved signals (Fig. 1C), indicating that these NTE residues retain high local mobility, without the severe conformational restriction expected for residues stably embedded in a dense crosslinked network. Conversely, in the CP-based ¹H-¹⁵N HETCOR experiment, strong signals were detected exclusively from the CSD, while CD signals were entirely absent (Fig. 1B). This indicates that the CSD is immobilized within large oligomeric assemblies with severely restricted mobility, whereas the CD retains relative motional freedom, making it in the intermediate regime invisible to both CP and INEPT spectra. Moreover, the observation of multiple chemical shifts for several CSD residues (e.g., R119, K125, M137, W142; Fig. 1B) suggests the CSD adopts polymorphic oligomeric states within the condensate. Together, these observations reveal a dynamic hierarchy within the condensates: the high mobility of phosphorylated NTE sites challenges their proposed role as stable multivalent hubs, while possible polymorphic CSD oligomers are consistent with the previously determined oligomeric structure by crystallography [12].

### NTE phosphorylation promotes phase separation by releasing NTE from the chromodomain

To assess phosphorylation-induced conformation changes in HP1α, we collected solution-state ¹H-¹⁵N HSQC spectra of phosphorylated (pHP1α) and its non-phosphorylated counterpart in the dispersed, non-condensed state. CSP analysis revealed that phosphorylation-induced changes localized predominantly to the NTE and CD (Fig. 2A). The NTE perturbations are readily attributable to the phosphorylation event itself; however, the CD chemical shift changes suggest that the NTE and CD may engage in direct intramolecular contact, which is perturbed upon phosphorylation.

**Fig. 2.**
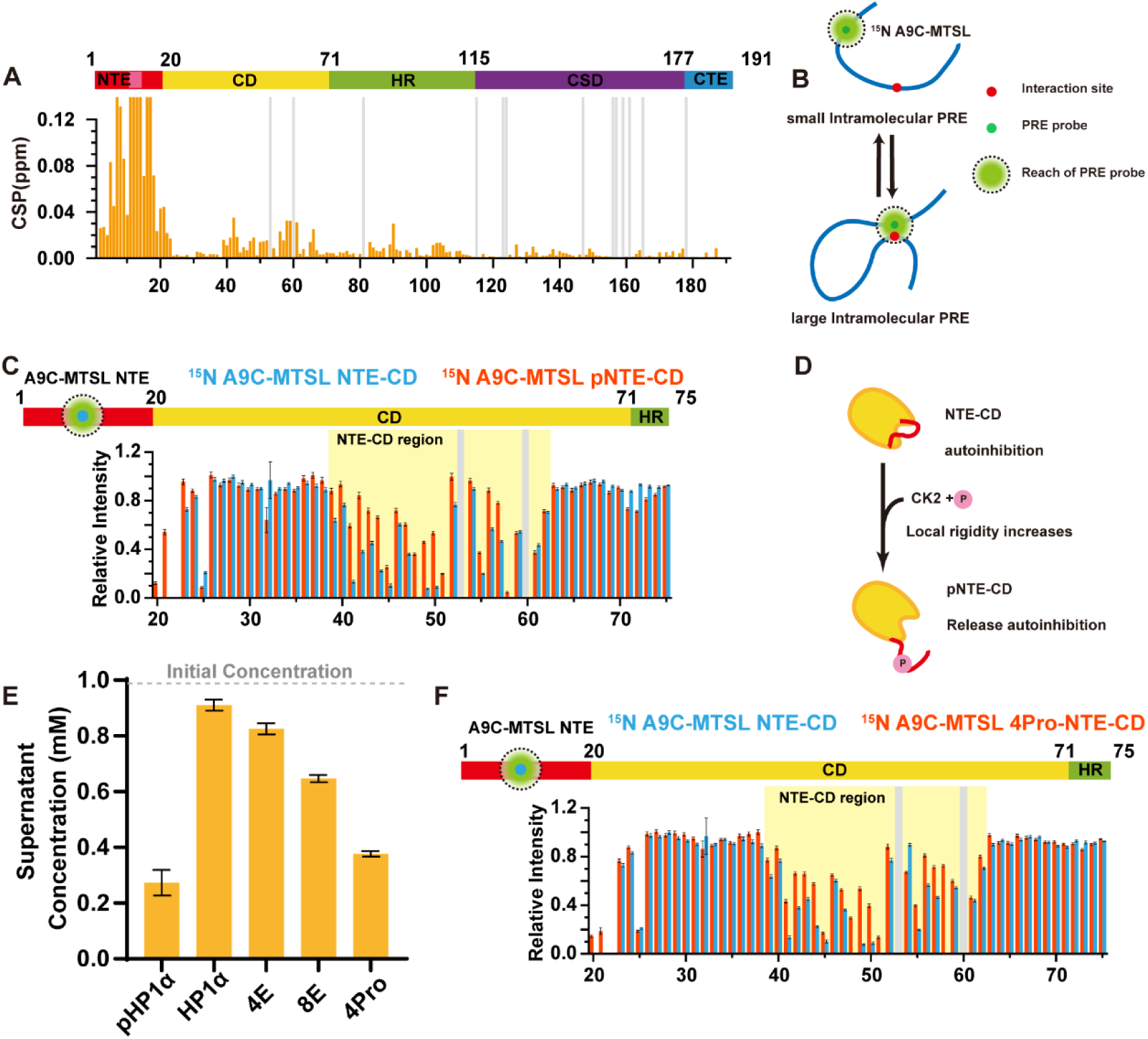
NTE phosphorylation promotes phase separation by relieving intramolecular autoinhibition. **(A)** CSP analysis of pHP1α versus non-phosphorylated HP1α. Phosphorylation-induced perturbations localize predominantly to the NTE and CD. **(B)** Intramolecular PRE schematic: A9C-MTSL spin label on the NTE probes intramolecular NTE-CD proximity. **(C)** PRE intensity ratios (Ipara/Idia) of NTE-CD (blue) and pNTE-CD (red). Non-phosphorylated NTE-CD shows pronounced attenuation across CD residues 39-62 (the NTE-contacting region), demonstrating close intramolecular contact; phosphorylation markedly reduces attenuation, indicating NTE dissociation from the CD. **(D)** Model: In the apo state, the NTE binds the CD, maintaining autoinhibition; phosphorylation drives NTE dissociation, relieving the constraint. **(E)** In vitro LLPS of full-length NTE mutants. Lower supernatant concentration indicates stronger phase separation. 4Pro promotes phase separation as effectively as true phosphorylation; 8E weakly promotes LLPS, whereas 4E is ineffective. **(F)** 4Pro-NTE-CD PRE recapitulates the pNTE-CD profile, confirming that increased local rigidity alone suffices to drive NTE dissociation from the CD.

To test this hypothesis, we used paramagnetic relaxation enhancement (PRE) to probe the spatial proximity between the NTE and the CD. A site-directed MTSL spin label attenuates signals within ∼25 Å, making PRE well suited for detecting transient, weak intramolecular contacts that are difficult to resolve by CSP alone [45-47]. We constructed an NTE-CD fragment bearing an A9C-MTSL spin label on the NTE (Fig. 2B). In the non-phosphorylated state, PRE revealed pronounced signal attenuation across CD residues 39-62 (the NTE-contacting region; Fig. 2C), directly demonstrating that the NTE is positioned in close proximity to the CD. Phosphorylation (pNTE-CD) markedly reduced this attenuation (Fig. 2C), indicating a phosphorylation-induced increase in the NTE-CD distance and disruption of their intramolecular contact. These observations establish that the NTE maintains HP1α in an autoinhibited conformation by binding the CD and occluding multivalent interaction interfaces; phosphorylation relieves this constraint by driving the NTE dissociation from the CD (Fig. 2D). This dissociation thus constitutes a critical conformational switch that relieves autoinhibition and initiates phase separation.

The phosphomimetic 4E mutant (SSSS→EEEE), widely used previously, fails to promote phase separation [16]. One explanation is insufficient charge compensation: four glutamates deliver far fewer negative charges than four phosphate groups. However, phosphorylation has also been reported to enhance local conformational rigidity [48], raising an alternative possibility—that rigidity, rather than charge, is the primary driver of phase separation. To distinguish between these possibilities, we constructed two NTE mutants: 4Pro (SSSS→PPPP), which restricts Cα-N bond rotation to increase rigidity without altering net charge; and 8E (SSSS→EEEEEEEE), which maximally compensates phosphorylation-associated negative charge. In vitro phase separation assays showed that 8E only weakly promoted phase separation, whereas 4Pro strongly induced it, to an extent comparable to true phosphorylation (Fig. 2E, Fig. S2). This pattern was recapitulated in cells: 4Pro significantly enhanced condensate formation, while 4E and 8E did not (Fig. 3A, B). PRE analysis of 4Pro confirmed that mimicking rigidity alone suffices to promote NTE release from CD, recapitulating the phosphorylation effect (Fig. 2F). Collectively, these results establish that the primary mechanism by which NTE phosphorylation promotes HP1α phase separation is relieving autoinhibition through NTE-CD dissociation, with charge playing only a secondary role.

**Fig. 3.**
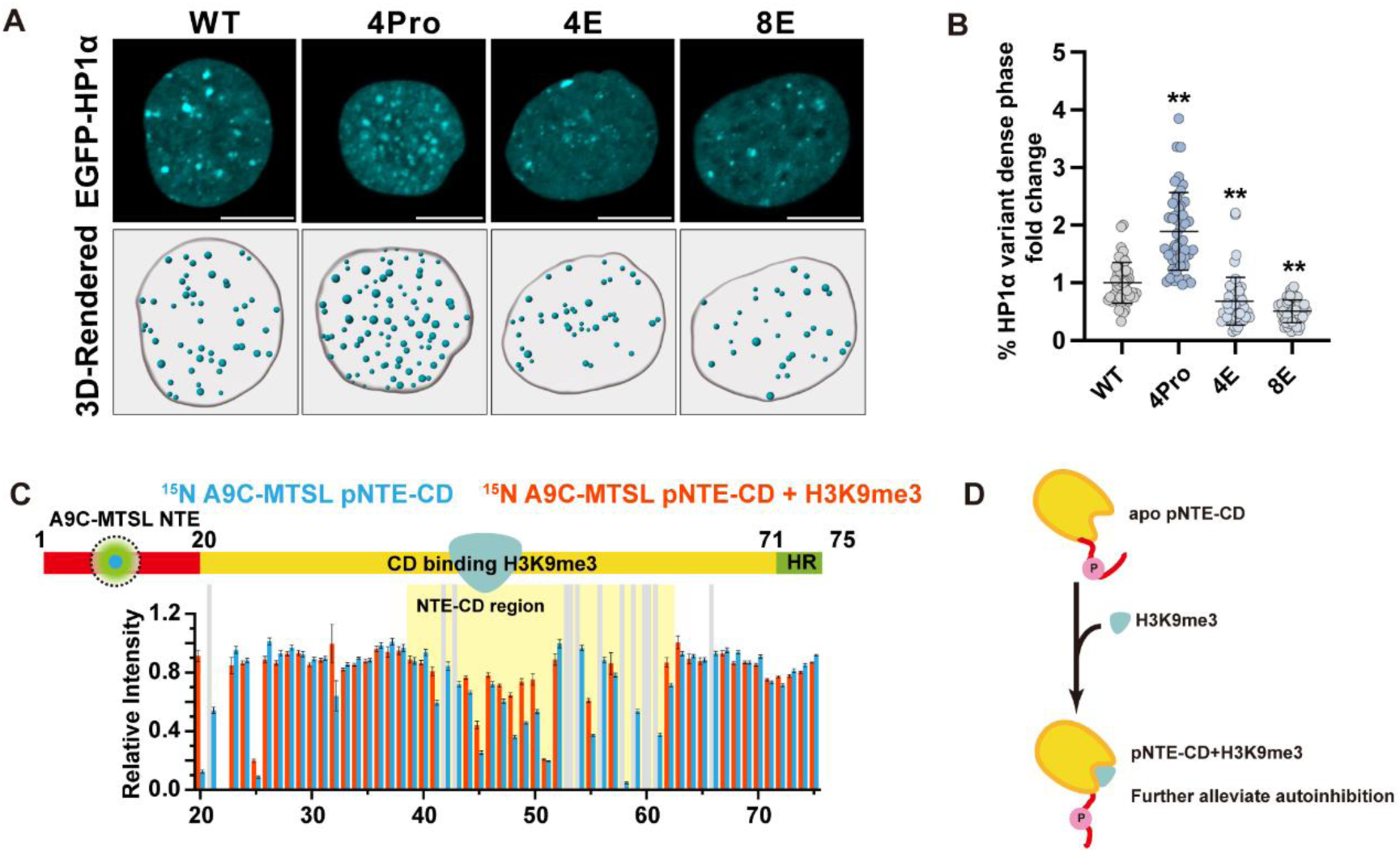
Cellular validation and allosteric modulation of NTE-CD autoinhibition by H3K9me3. (A) Live-cell imaging of EGFP-HP1α condensates in HEK293 cells expressing WT, 4Pro, 4E, or 8E NTE mutants. Scale bar: 10 µm. (B) Quantification of the ratio of condensed-phase HP1α fluorescence to total nuclear HP1α fluorescence, normalized to the WT control (set to 1). 4Pro significantly increases this ratio, whereas 4E and 8E significantly decrease it (**p < 0.01 versus WT). Horizontal lines denote the mean, and error bars indicate the standard deviation, n = 50 nuclei. Data are the mean ± SD. The P value was calculated by one-way ANOVA. (C) PRE intensity ratios (I_para_/I_dia_) of A9C-MTSL pNTE-CD in the absence (blue) or presence (orange) of H3K9me3. H3K9me3 further reduces PRE attenuation across the CD region, indicating enhanced NTE dissociation from the CD. (D) Model: H3K9me3 binding to the CD allosterically enhances NTE dissociation, synergizing with phosphorylation to relieve autoinhibition.

Previous work showed that H3K9me3 promotes pHP1α phase separation [16]. Notably, NMR titration indicates that H3K9me3 binds the CD at the same interface engaged by the NTE [48], suggesting that H3K9me3 may directly displace or allosterically release the NTE. To test this, we titrated H3K9me3 into pNTE-CD and monitored PRE attenuation. H3K9me3 markedly reduced PRE attenuation across the NTE-contacting region of the CD (residues 39-62) (Fig. 3C), directly demonstrating enhanced NTE dissociation from the CD (Fig. 3D). This provides a mechanistic basis for the synergistic promotion of phase separation by H3K9me3.

In summary, these results revise the prevailing charge-centric model by establishing that NTE phosphorylation promotes HP1α phase separation primarily through increased local rigidity and release of the NTE from the CD, thereby relieving intramolecular autoinhibition. Electrostatic charge contributes only secondarily. This conformational release mechanism is further supported by H3K9me3, which binds the CD and similarly enhances NTE dissociation. Together, these findings provide a mechanistic basis for the coordinated regulation of HP1α condensation by phosphorylation and H3K9me3.

### CTE suppresses phase separation by blocking the HR-CSD interaction

Previous studies proposed that CTE suppresses HP1α phase separation by competing with the NTE for HR binding [16, 20, 26, 28]. However, whether CTE additionally modulates HR interactions with other domains to suppress phase separation remains unclear. To directly assess this, we constructed two truncated fragments: HR-CSD and HR-CSD-CTE. At high protein concentrations, solution-state ¹H-¹⁵N HSQC spectra of HR-CSD showed near-complete disappearance of CSD peaks and pronounced line broadening of HR signals (Fig. 4A), indicating extensive aggregation with markedly accelerated transverse relaxation. In contrast, HR-CSD-CTE showed neither peak loss nor line broadening (Fig. 4A), indicating that CTE suppresses higher-order assembly, maintaining the protein in a monodispersed state under conditions where HR-CSD already aggregates. In vitro phase separation assays confirmed that both fragments undergo LLPS, but HR-CSD exhibited a markedly lower concentration threshold than HR-CSD-CTE (Fig. 5D, Fig. S3).

**Fig. 4.**
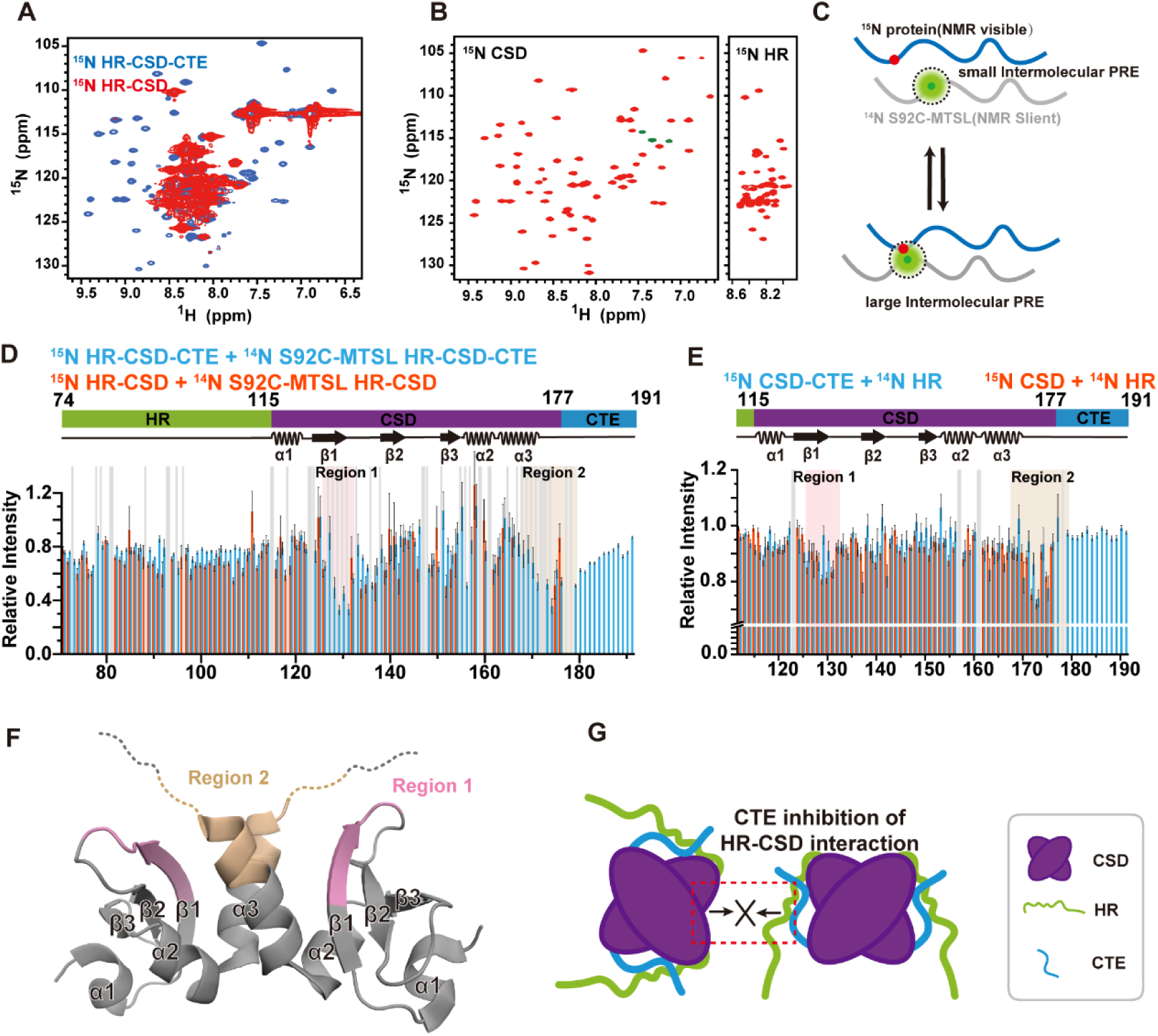
HR-CSD interaction drives phase separation and is antagonized by CTE. **(A)** Solution-state ¹H-¹⁵N HSQC spectra of HR-CSD versus HR-CSD-CTE. HR-CSD shows near-complete disappearance of CSD peaks due to aggregation, whereas HR-CSD-CTE retains peaks. **(B)** Isolated CSD and HR, at high concentration, both yielded highly resolved ¹H-¹⁵N HSQC peaks, indicating that neither undergoes extensive aggregation independently. **(C)** PRE schematic for intermolecular HR-CSD mapping. **(D)** Intermolecular PRE (S92C-MTSL in HR). In HR-CSD, strong spontaneous aggregation causes severe line broadening and signal loss of CSD peaks. In contrast, HR-CSD-CTE yields much better spectral quality, revealing specific contacts at CSD Region 1 (residues 126-132) and Region 2 (residues 168-179). **(E)** Titration of natural abundance HR into ¹⁵N-labeled CSD or ¹⁵N-labeled CSD-CTE. HR-induced CSPs in Region 1 and 2 are weakened by CTE. **(F)** Mapping of HR binding Region 1 and 2 on the CSD structure (PDB: 9CEA). **(G)** Model: CTE blocks HR-CSD interaction.

**Fig. 5.**
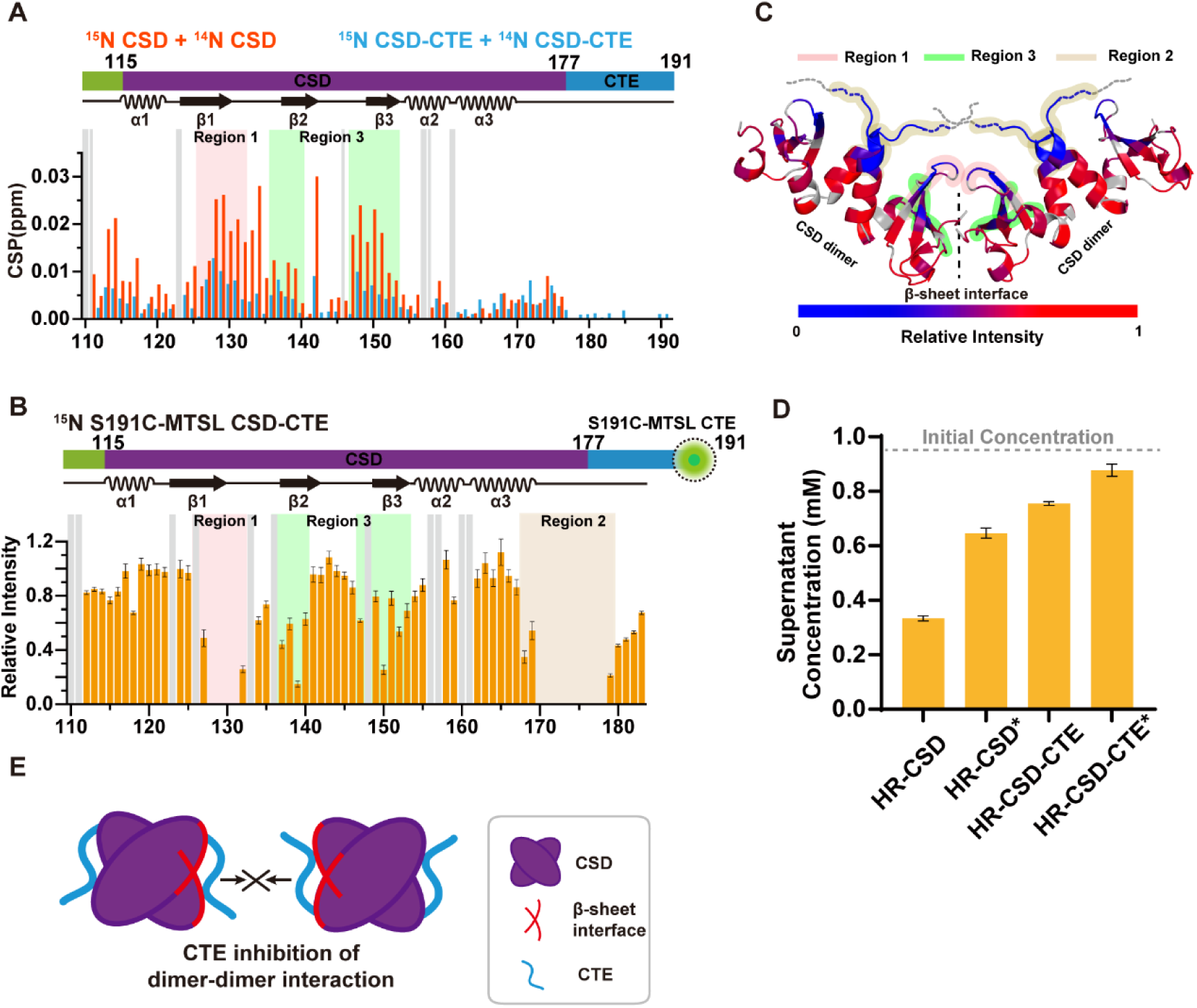
CTE directly binds CSD to block dimer-dimer interactions and higher-order assembly. **(A)** CSP analysis of ¹⁵N-CSD upon titration of natural-abundance CSD (blue) or natural-abundance CSD-CTE (orange). Inter-dimer contacts are markedly suppressed in the presence of CTE. **(B)** Intramolecular PRE of S191C-MTSL-labeled CTE in the CSD-CTE fragment. Attenuation maps to Region 1, 2, and 3 on the CSD. **(C)** Mapping of PRE-defined Region 3 onto the CSD crystal structure (PDB: 9CDW), revealing its location at the core of the β-sheet interface that mediates higher-order CSD oligomerization. **(D)** Supernatant protein concentrations after centrifugation of HR-CSD, HR-CSD-CTE, and their L139E/L150E mutants (HR-CSD* and HR-CSD-CTE*). Lower supernatant concentration indicates stronger phase separation. **(E)** Model: CTE directly binds the CSD β-sheet interface to block interaction between dimers.

To test whether HR or CSD alone drives assembly, we purified both domains individually. At high concentrations, both yielded high-resolution HSQC spectra with strong signals (Fig. 4B), indicating that neither forms aggregates on its own. Thus, HR-CSD cooperation is required for higher-order assembly. In line with this, HR and CSD do not undergo phase separation on their own (Fig. S3).

To probe intermolecular interactions, we employed an isotope-filtered PRE scheme: the MTSL spin label was introduced on a ¹⁴N-labeled, NMR-invisible protein, while signal attenuation was monitored in its ¹⁵N-labeled partner, such that any observed attenuation arises exclusively from intermolecular contacts (Fig. 4C). We introduced S92C-MTSL in HR and assessed intermolecular PRE against ¹⁵N-labeled HR-CSD and HR-CSD-CTE. In HR-CSD, strong spontaneous aggregation caused severe line broadening and signal loss of CSD peaks (Fig. 4D). In HR-CSD-CTE, where aggregation is suppressed, CSD signals were retained with sufficient quality for PRE analysis. The PRE profile revealed direct HR-CSD interactions primarily localized to Region 1 (residues 126-132) and Region 2 (residues 168-179) (Fig. 4D, F). Notably, these contacts remain detectable in the presence of CTE, indicating that CTE attenuates the HR–CSD interaction rather than abolishing it.

To further characterize HR-CSD binding and CTE regulation, we titrated unlabeled HR into ¹⁵N CSD or ¹⁵N CSD-CTE. HR-induced signal attenuations appeared at Region 1 and 2, corroborating the PRE-mapped interface. These attenuations were clearly reduced in the presence of CTE, demonstrating that CTE weakens the HR-CSD interaction (Fig. 4E).

### CTE directly binds CSD to inhibit the dimer-dimer interactions

Inter-dimer interactions between CSDs have been implicated in higher-order HP1α assembly [12], but whether CTE directly regulates them remains unclear. To test this, we titrated natural-abundance CSD into ¹⁵N-labeled CSD and natural-abundance CSD-CTE into ¹⁵N-labeled CSD-CTE, respectively. CSP analysis revealed pronounced perturbations in ¹⁵N CSD upon addition of natural-abundance CSD, indicating inter-dimer interactions; these perturbations were markedly suppressed in the ¹⁵N CSD-CTE sample (Fig. 5A), demonstrating that CTE inhibits CSD dimer-dimer contacts. To map the CTE-CSD interface responsible for this inhibition, we introduced S191C-MTSL in CTE and performed intramolecular PRE on CSD-CTE. MTSL strongly attenuated signals from Region 1 and 2 on the CSD, with additional attenuation in Region 3 (residues 136-140 and 147-153), identifying three direct contact regions (Fig. 5B, C).

The three CTE-contacted regions offer a structural explanation for this inhibition. First, analysis of the inter-dimer CSPs in Fig. 5A showed that they localize predominantly to Region 1 and 3, indicating that these regions mediate CSD dimer-dimer association. Independently, previous crystallographic work on the CSD-KAP1_HBox_ complex (PDB: 9CDW) captured a β-sheet-mediated packing interface between adjacent CSD dimers, based on which a higher-order CSD oligomerization model was constructed [12]. Mapping our PRE-defined regions onto this structure places Region 3 at the core of the β-sheet interface (Fig. 5C). The CTE-contacted regions thus overlap the inter-dimer interface, and CTE binding sterically occludes it, accounting for the suppressed inter-dimer contacts observed in the CSD-CTE titration. This experimentally defined mechanism is consistent with MD simulations proposing that the CTE modulates the CSD oligomerization interface [13].

Previous studies identified the hydrophobic residues L139 and L150 within Region 3 as key mediators of CSD oligomerization [13]; the L139E/L150E double mutant disrupts CSD oligomerization and impairs HP1α phase separation [12-14]. We introduced this mutation into HR-CSD and HR-CSD-CTE (designated HR-CSD* and HR-CSD-CTE*, respectively) and observed markedly reduced phase separation in both constructs (Fig. 5D), confirming that the β-sheet-mediated dimer-dimer interface is essential for LLPS. Notably, CTE retained its inhibitory effect in the mutant background (HR-CSD-CTE* versus HR-CSD*). Because the HR-binding sites on the CSD (Region 1 and 2) are spared by the L139E/L150E mutation, this residual suppression is most readily explained by CTE’s blockade of the HR-CSD interaction established in Fig. 4, which remains intact in the mutant. These comparisons therefore resolve CTE’s inhibitory function into two mechanistically separable components.

Together, these results demonstrate that CTE suppresses HP1α phase separation through a second, distinct mechanism: direct binding to the CSD β-sheet interface, particularly Region 1 and 3, which mediate CSD dimer interaction. Combined with the HR-CSD interaction blockade established in Fig. 4, these findings reveal that CTE employs dual inhibition to restrain HP1α condensation: by weakening intermolecular crosslinking between HR and CSD, and by directly occluding the CSD dimer interaction interface (Fig. 4G, 5E).

### Poly-PR exerts a concentration-dependent biphasic effect on pHP1α phase separation

To mimic disease-associated poly-PR repeats, we used PR20 (20 repeats of poly-PR) as a model peptide in vitro [37, 39]. Systematically varying the concentrations of both pHP1α and PR20 across a two-dimensional grid yielded a concentration-dependent phase diagram in which phase separation was confined to a defined PR20 concentration window, emerging at the lower boundary and dissolving beyond the upper boundary (Fig. 6A). To quantitatively validate this trend, we measured supernatant pHP1α at a fixed concentration of 150 µM across varying PR20 concentrations. Below 350 µM, PR20 promoted phase separation, as evidenced by reduced supernatant pHP1α (Fig. 6B, Fig. S4A). Notably, phase separation occurred at pHP1α concentrations below the threshold of pHP1α alone, indicating that poly-PR shifts the phase boundary of the mixture toward lower pHP1α concentrations. Above 350 µM, PR20 reversed this effect, suppressing and ultimately abolishing condensation at higher concentrations, as evidenced by increased supernatant pHP1α (Fig. 6A, B). PA20 (20 repeats of poly-PA) showed no effect at any concentration tested (Fig. 6B), confirming that the biphasic regulation is specific to the PR repeat, consistent with the distinct pathogenic activities of PR and PA dipeptide repeats in C9orf72 disease [35, 49].

**Fig. 6.**
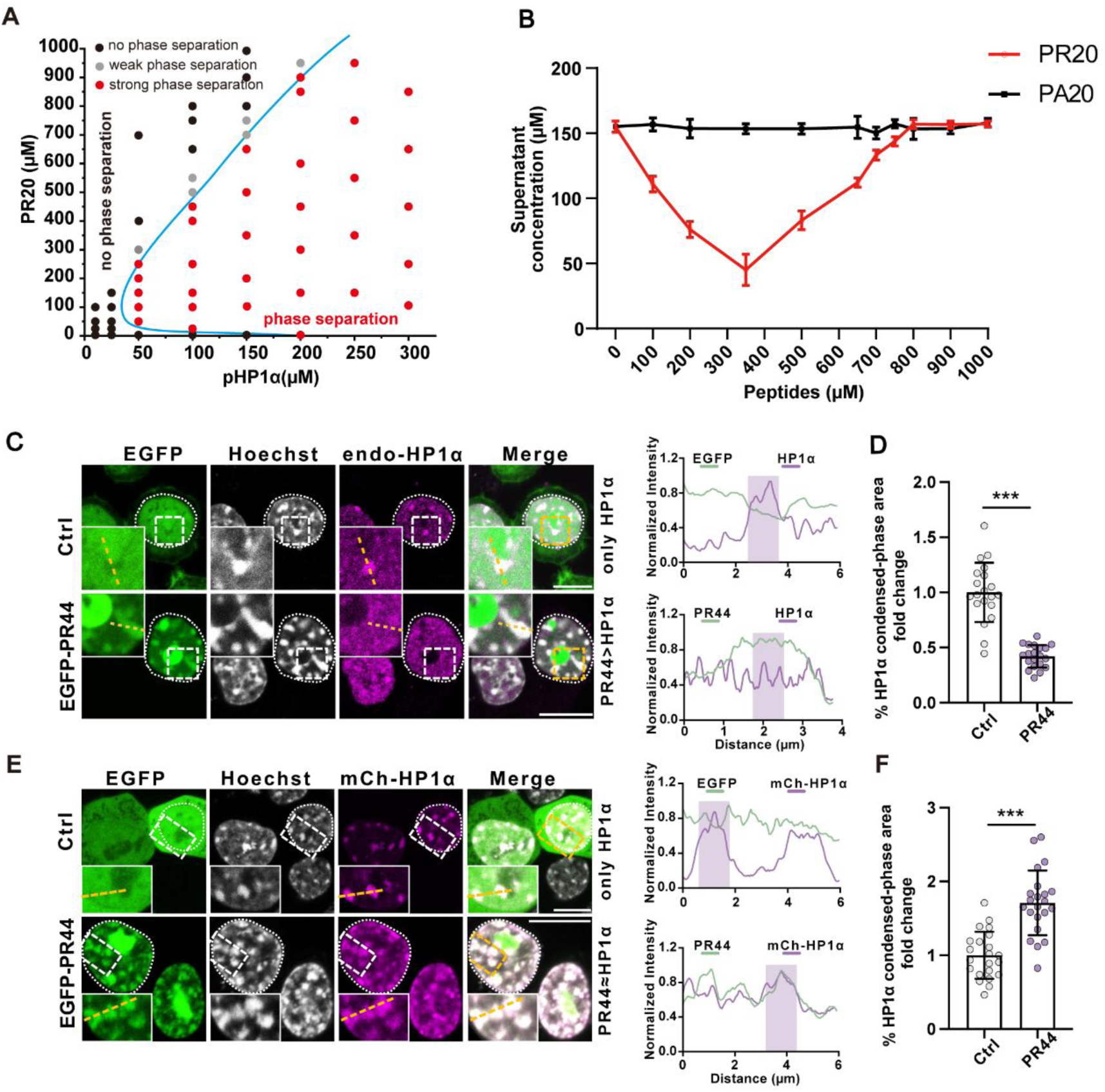
Poly-PR exerts a concentration-dependent biphasic effect on pHP1α phase separation in vitro and in cells. **(A)** Phase diagram of pHP1α versus PR20. Red, strong LLPS; gray, weak LLPS; black, no LLPS. **(B)** Supernatant pHP1α concentration after centrifugation at fixed 150 µM pHP1α across varying PR20 concentrations. Lower supernatant concentration indicates stronger phase separation. PR20 promotes LLPS at <350 µM and inhibits LLPS at >350 µM; PA20 is inactive. **(C)** Representative immunofluorescence images of endogenous HP1α in N2A cells overexpressing EGFP-PR44, shifting the poly-PR: HP1α ratio toward the inhibition regime. Poly-PR disperses endogenous HP1α condensates. Dashed lines indicate scan paths; the corresponding normalized fluorescence intensity profiles (EGFP, HP1α) are shown on the right. Scale bar, 10 µm. **(D)** Quantification of the condensed-phase fraction of HP1α for the experiments shown in (C), calculated as the ratio of condensate fluorescence area to total nuclear fluorescence area and normalized to the control. Data are mean ± SD; unpaired t-tests. n = 20 nuclei, ***p < 0.001. **(E)** Representative live-cell images of N2A cells co-overexpressing mCherry-HP1α and EGFP-PR44, shifting the poly-PR: HP1α ratio toward the promotion regime. HP1α and PR44 undergo co-LLPS. Yellow dashed lines indicate scan paths; the corresponding normalized fluorescence intensity profiles of EGFP and mCherry are shown on the right. Scale bar: 10 µm. **(F)** Quantification of the condensed-phase fraction of HP1α for the experiments shown in (E), calculated as the ratio of condensate fluorescence area to total nuclear fluorescence area and normalized to the control. Data are mean ± SD; unpaired t-tests. n = 22 nuclei, ***p < 0.001.

To test whether the concentration-dependent biphasic regulation operates in cells, we set up two poly-PR: HP1α stoichiometric regimes against the same overexpressed poly-PR background. In the first regime, EGFP-PR44 was overexpressed alone; with endogenous HP1α as the readout, the low-abundance HP1α pool places the poly-PR: HP1α ratio in the saturation regime. Immunofluorescence showed that endogenous HP1α condensates were markedly dispersed (Fig. 6C, D), mirroring the high-ratio dissolution observed in vitro. In the second regime, mCherry-HP1α was co-overexpressed with EGFP-PR44, increasing the total HP1α pool and, by design, lowering the poly-PR: HP1α ratio toward the promotion regime. Live-cell imaging revealed restored condensate formation, with HP1α and PR44 undergoing co-condensation (Fig. 6E, F). Thus, with poly-PR overexpressed in both regimes, lowering the poly-PR: HP1α ratio converted the outcome from condensate dissolution to co-condensation. These cellular observations recapitulate both regimes of the in vitro biphasic regulation, demonstrating that the fate of HP1α condensates is dictated by the poly-PR: HP1α stoichiometry rather than by the mere presence of poly-PR.

Extending these findings to a pathological setting, we expressed EGFP-PR30 in neonatal mouse brain by intracerebroventricular AAV injection, a context in which the AAV-expressed PR30 places the ratio in the saturation regime relative to endogenous HP1α. Accordingly, PR30-expressing neurons showed diffuse endogenous HP1α with significantly suppressed puncta formation compared to neighboring non-expressing cells (Fig. S4B), consistent with condensate dissolution in cells.

### Poly-PR binds the NTE to modulate pHP1α interaction networks

The stoichiometry dependence of this regulation argues against simple competitive inhibition and instead implicates a multivalent interaction mode. To elucidate the molecular basis of this biphasic regulation, we titrated PR20 into pHP1α and monitored binding by NMR. Spectra were recorded at 289 K and 308 K, temperatures chosen to optimize signal detection for flexible IDRs and folded domains, respectively. CSP analysis revealed that PR20 binding primarily affected the NTE and CD, with minimal perturbations elsewhere (Fig. 7A). Notably, the CSP patterns in the CD closely resembled those induced by NTE phosphorylation (Fig. 2A), suggesting that PR20 binding perturbs the NTE-CD intramolecular interface in a manner analogous to phosphorylation.

**Fig. 7.**
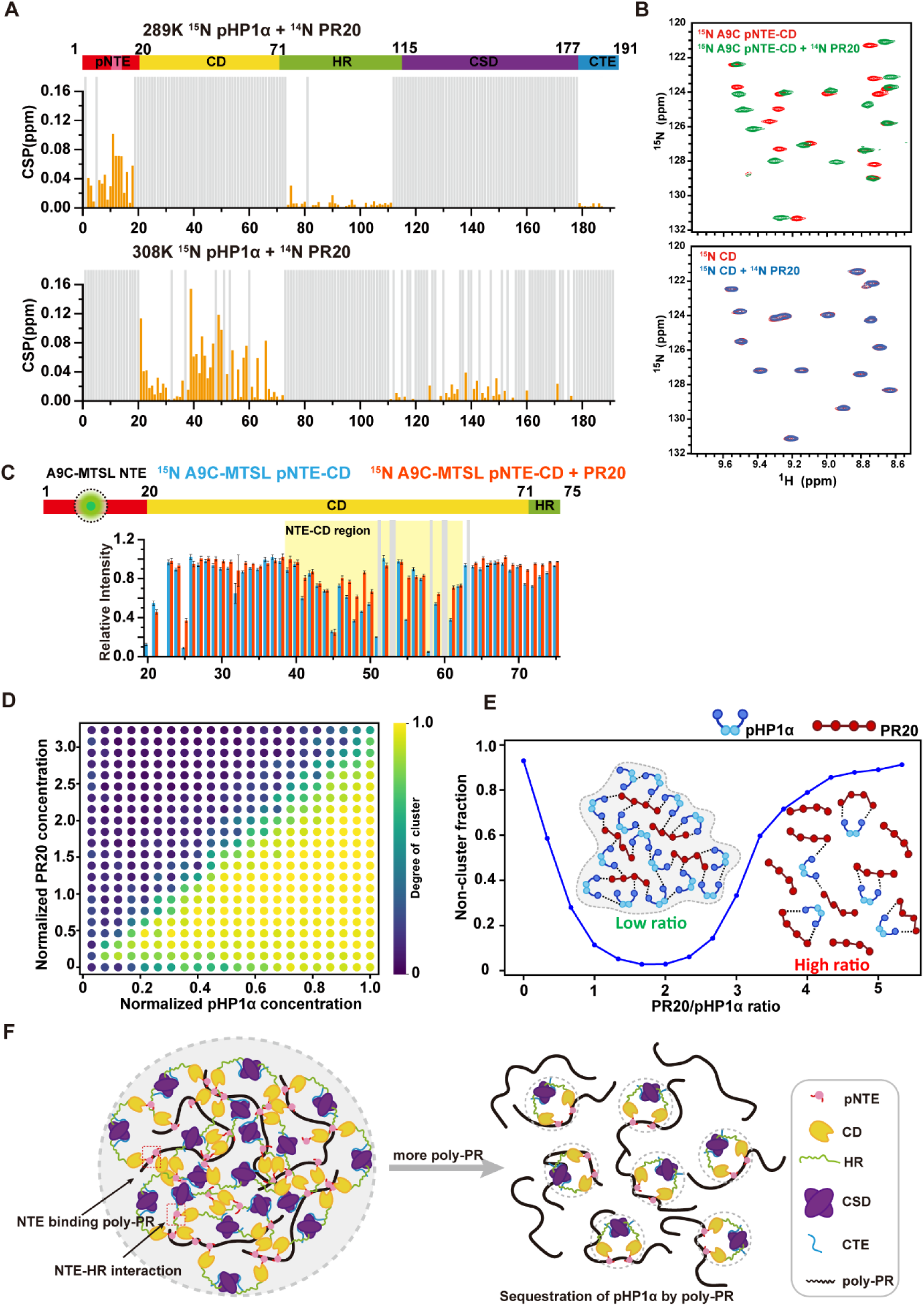
Poly-PR binds the NTE to relieve autoinhibition and exerts biphasic regulation via multivalent crosslinking and site saturation. **(A)** CSP analysis of pHP1α upon PR20 titration. Spectra were recorded at 289 K and 308 K to optimize signal detection for flexible IDRs and folded domains, respectively. Perturbations are predominantly related to the NTE and CD. **(B)** PR20 titration into isolated CD (no CSP) versus pNTE-CD (strong CSP), establishing direct NTE binding. **(C)** PRE of A9C-MTSL pNTE-CD in the absence (blue) or presence (red) of PR20. PR20 reduces PRE attenuation across the NTE-contacting region, indicating NTE dissociation from the CD. **(D)** Monte Carlo simulation phase diagram of pHP1α-PR20 mixture across varying PR20:pHP1α stoichiometries. **(E)** Simulated non-cluster fraction versus PR20:pHP1α ratio. Low ratios promote large interconnected clusters, indicative of enhanced phase separation; high ratios fragment the network into isolated small aggregates, indicative of condensate dissolution. **(F)** Model: At low concentrations, multivalent PR20 engages multiple NTEs to crosslink pHP1α molecules, promoting phase separation. At high concentrations, NTE sites become saturated, isolating pHP1α into 1:1 complexes and dissolving condensates.

To identify the direct binding site of PR20, we titrated PR20 into isolated CD and pNTE-CD fragments. PR20 induced no detectable CSP on the isolated CD, ruling it out as the direct binding interface (Fig. 7B). In contrast, the pNTE-CD fragment showed pronounced CSP changes, indicating that PR20 binds directly to the NTE rather than the CD. The CD CSPs arise indirectly following NTE binding.

The indirect CD perturbations suggest that PR20 binding alters the intramolecular NTE-CD contact. To test this directly, we applied the intramolecular PRE readout established in Fig. 2, in which an A9C-MTSL label on the NTE reports its proximity to the CD through attenuation across CD residues 39-62. Phosphorylation alone already reduces this attenuation relative to the non-phosphorylated state (Fig. 2C). Addition of PR20 to pNTE-CD reduced attenuation across this region relative to pNTE-CD alone (Fig. 7C), indicating that PR20 binding displaces the NTE from the CD beyond the release achieved by phosphorylation. This conformational consequence corroborates direct NTE engagement by PR20. However, NTE release alone cannot account for the concentration-dependent biphasic regulation.

To elucidate the mechanism underlying this biphasic behavior, we performed Monte Carlo simulations of multivalent binding across varying PR20:pHP1α stoichiometries (Fig. S5). In the model, each pHP1α presents one PR20-binding site on its NTE, and each flexible PR20 chain can engage up to four NTEs simultaneously. Cluster formation was evaluated from the connectivity of the simulated binding network and summarized as the degree of clustering across normalized pHP1α and PR20 concentrations. The resulting phase diagram revealed a biphasic dependence on pHP1α:PR20 amount ratios: increasing PR20 initially enhanced clustering, whereas excessive PR20 reduced cluster formation despite the continued presence of both components (Fig. 7D). At low ratios, simulations predicted expanded, highly interconnected clusters in which individual PR20 chains bridge multiple pHP1α molecules (Fig. 7D, E), recapitulating the experimentally observed shift of the phase boundary toward lower pHP1α concentrations. At high ratios, this trend inverted sharply: as NTE sites approached saturation, each NTE became occupied by a single PR20 chain, forming 1:1 NTE-PR20 complexes that abolish multivalent bridging; consequently, continuous macro-clusters fragmented into isolated small aggregates, and inter-protein connectivity was lost, as reflected by the rising non-cluster fraction (Fig. 7D, E). The simulated phase diagram thus reproduces the full biphasic profile observed experimentally (Fig. 7D; compare Fig. 6A), supporting a model in which stoichiometry-dependent PR20-mediated crosslinking and subsequent binding-site saturation are sufficient to account for the biphasic regulation.

Based on these dynamic changes, we propose a concentration-dependent interaction model (Fig. 7F): At low concentrations, highly cationic poly-PR binds electrostatically to the negatively charged NTE, stripping it from CD and relieving autoinhibition. Meanwhile, a single poly-PR chain can simultaneously engage multiple NTE motifs, acting as a molecular adhesive tape that crosslinks several pHP1α molecules. This multivalent bridging lowers the critical concentration for phase separation and promotes droplet formation. As poly-PR concentration rises, NTE sites become progressively saturated; in the limiting case, each NTE is occupied by one poly-PR molecule in a 1:1 complex. Such 1:1 occupancy abolishes multivalent bridging, so that pHP1α can no longer assemble into a crosslinked network, and phase separation is completely inhibited.

## Discussion

HP1α is critical for heterochromatin formation and maintenance [1-4]. However, the classical IDR-driven LLPS model fails to account for several key experimental observations in this study, suggesting that multivalent IDR interactions are not the sole driver of phase separation and that folded domains play a more active regulatory role than previously appreciated. By integrating biochemical and NMR approaches, we systematically demonstrate that HP1α phase separation relies not only on IDR interactions but crucially on cooperative regulation between IDRs and folded domains. We further show that ALS-associated poly-PR engages this network at the NTE, exerting stoichiometry-dependent biphasic control over HP1α condensation: multivalent bridging promotes condensation at low poly-PR: HP1α ratios, whereas site saturation dissolves condensates at high ratios. This work reframes the molecular model of HP1α phase separation from an IDR-centric to a whole-molecule cooperative perspective, establishes folded domains as key regulators in multi-domain protein phase separation, and illustrates how a pathogenic dipeptide repeat can exploit a condensate’s interaction network, providing a molecular framework for understanding heterochromatin disruption in C9orf72-linked disease.

The conventional view attributes NTE phosphorylation-driven phase separation to increased negative charge and enhanced electrostatic interactions with HR [16, 17, 20, 26, 28]. Our data revise this charge-centric model: phosphorylation acts primarily by increasing the local rigidity of the NTE, which drives its release from the CD and exposes interaction interfaces, with charge contributing only secondarily. This reinterpretation resolves a long-standing puzzle in the field: phosphomimetic mutants such as 4E fail to promote phase separation not because charge compensation is insufficient, but because glutamate substitutions mimic electrostatics without the conformational constraint imposed by phosphate groups. More broadly, this distinction cautions against the default interpretation of phosphomimetic phenotypes, which capture charge but not necessarily the structural consequences of phosphorylation. The same release mechanism is engaged by H3K9me3, indicating that phosphorylation and the histone mark converge on a common autoinhibitory switch, thereby coupling HP1α condensation directly to chromatin context.

Previous studies have largely overlooked the active role of CSD in phase separation, while HR has been viewed merely as a competitive NTE/CTE-binding region or a nonspecific DNA-binding site [16, 18, 28]. Our findings reposition both elements: direct HR-CSD interaction provides a driving force for condensation, while the CTE acts as a dual gatekeeper, occluding both the HR–CSD interface and the β-sheet interface that mediates CSD dimer–dimer contacts. Folded-domain surfaces thus participate actively in the regulatory logic of condensation rather than serving as passive scaffolds. Notably, our experimentally defined interfaces agree with predictions from simulations based on truncated constructs lacking the flexible regions [13], providing long-sought experimental support for those models and suggesting that their conclusions extend to the full-length protein. These findings may also help explain the diverse effects of CSD ligands on HP1α phase separation: PXVXL-motif proteins bind the CSD dimer but exert markedly divergent effects, generally attributed to their charge properties [16, 28]. Whether these ligands act by modulating the CSD interaction interfaces defined here, rather than through charge alone, remains to be investigated.

Solid-state NMR of the condensate at residue resolution reveals a pronounced dynamic hierarchy (Fig. 1B, C). Strong CP signals arise exclusively from the CSD, identifying it as a rigid, motion-restricted assembly, whereas the CD, invisible in both CP and INEPT spectra, occupies an intermediate dynamic regime, and the NTE phosphorylation sites retain high local mobility. The severe motional restriction of the CSD indicates that it assembles into large oligomers within the condensate, and the multiple chemical shifts observed for several CSD residues further suggest that these assemblies are polymorphic, consistent with crystal structures showing that the CSD β-sheet interface supports multiple oligomeric forms (e.g., PDB 3I3C, 9CEA) [12]. Solid-state NMR thus provides direct access to the structure, dynamics, and interactions within the condensed phase, complementing solution-state characterization of the dispersed state; this in situ capability is essential for a complete understanding of condensate architecture [50, 51].

Poly-PR interacts with multiple proteins and causes diverse functional perturbations [34, 35]. Our finding that it regulates HP1α condensation biphasically, through stoichiometry-dependent crosslinking and saturation, adds a mechanistic dimension to this picture: the outcome of poly-PR exposure is dictated by the poly-PR: HP1α ratio rather than by the mere presence of the peptide. At the same time, although PR20 promotes HP1α droplet formation within a defined concentration window, whether the resulting PR–HP1α co-condensates are biophysically and functionally equivalent to HP1α-only condensates remains unclear; the promoted state may not be functional. Similar biphasic regulation has been reported for nucleophosmin (NPM1) and p53 [39, 52], suggesting a common mechanism by which DPRs perturb condensates: at low stoichiometry they crosslink client proteins and shift phase boundaries, whereas at high stoichiometry they saturate client binding sites and sequester them into assembly-incompetent complexes (Fig. 7F). Whether the cooperative interaction defined here is universal across HP1 isoforms or has diverged with their less conserved IDRs remains an open question.

## Acknowledgment

This work was supported by grants from the National Key R&D Program (2025YFA1308801), CAS Project for Young Scientists in Basic Research, Grant No. YSBR-068, the National Natural Science Foundation of China (32090040, 31971128, 22533005), Fundamental and Interdisciplinary Disciplines Breakthrough Plan of the Ministry of Education of China (JYB2025XDXM505), Guizhou Provincial Science and Technology Projects (QIANKEHEPINGTAI KXJZ [2024] 030).

## Author Contributions Statement

YZ.Z., S.D., H.L., T.S., Z.H., Y.Z., and S.X. conceived and designed the experiments. YZ.Z. performed most of the biochemical and NMR experiments. S.D. performed the cell and animal experiments. H.L. carried out the simulations. J.L. and C.S. helped with the NMR experiments. YZ.Z., S.X., Y.Z., Z.H. and T.S. wrote the manuscript.

## Reference

1. James, T.C. and S.C.R. Elgin, Identification of a Nonhistone Chromosomal Protein Associated with Heterochromatin in Drosophila-Melanogaster and Its Gene. Molecular and Cellular Biology, 1986. 6(11): p. 3862–3872.

2. Lachner, M., et al., Methylation of histone H3 lysine 9 creates a binding site for HP1 proteins. Nature, 2001. 410(6824): p. 116-120.

3. Maison, C. and G. Almouzni, HP1 and the dynamics of heterochromatin maintenance. Nature Reviews Molecular Cell Biology, 2004. 5(4): p. 296–304.

4. Zhu, Y., et al., HP1 enhances radio-and chemotherapy resistance in osteosarcoma via PARP1-dependent transcriptional repression and HR repair. Cancer Letters, 2026. 639.

5. Bannister, A.J., et al., Selective recognition of methylated lysine 9 on histone H3 by the HP1 chromo domain. Nature, 2001. 410(6824): p. 120-124.

6. Nishibuchi, G., et al., N-terminal phosphorylation of HP1α increases its nucleosome-binding specificity. Nucleic Acids Research, 2014. 42(20): p. 12498–12511.

7. Jacobs, S.A., et al., Specificity of the HP1 chromo domain for the methylated N-terminus of histone H3. Embo Journal, 2001. 20(18): p. 5232–5241.

8. Strom, A.R., et al., HP1alpha is a chromatin crosslinker that controls nuclear and mitotic chromosome mechanics. Elife, 2021. 10.

9. Richart, A.N., et al., Characterization of chromoshadow domain-mediated binding of heterochromatin protein 1alpha (HP1alpha) to histone H3. J Biol Chem, 2012. 287(22): p. 18730–7.

10. Mendez, D.L., et al., The HP1a disordered C terminus and chromo shadow domain cooperate to select target peptide partners. Chembiochem, 2011. 12(7): p. 1084–96.

11. Colmenares, Serafin U., et al., Expanding the HP1a-binding consensus and molecular grammar for heterochromatin assembly. Nucleic Acids Research, 2025. 53(19).

12. Gaurav, N., et al., The HP1 box of KAP1 organizes HP1α for silencing of endogenous retroviral elements in embryonic stem cells. Nature Communications, 2025. 16(1).

13. Kumar, A., et al., Mechanistic Insights into HP1α CSD Oligomerization and the Role of PxVxL Motif-Containing Proteins. Journal of Chemical Information and Modeling, 2025. 65(16): p. 8768–8780.

14. Gaurav, N., et al., HP1γ self-assembles and cooperates with KAP1 in repression of long noncoding RNA AI662270 in ESCs. Cell Reports, 2026. 45(2).

15. Zhao, T. and J.C. Eissenberg, Phosphorylation of Heterochromatin Protein 1 by Casein Kinase II Is Required for Efficient Heterochromatin Binding inDrosophila. Journal of Biological Chemistry, 1999. 274(21): p. 15095–15100.

16. Larson, A.G., et al., Liquid droplet formation by HP1alpha suggests a role for phase separation in heterochromatin. Nature, 2017. 547(7662): p. 236-240.

17. Furukawa, A., et al., A dynamic structural unit of phase-separated heterochromatin protein 1α as revealed by integrative structural analyses. Nucleic Acids Research, 2025. 53(6).

18. Keenen, M.M., et al., HP1 proteins compact DNA into mechanically and positionally stable phase separated domains. Elife, 2021. 10.

19. Zhai, Z., et al., Revealing site-specific molecular features mediating contacts of heterochromatin protein 1 α (HP1α) and the interactions with chromatin and DNA in LLPS environments. Communications Biology, 2025. 8(1).

20. Phan, T.M., et al., Interplay between charge distribution and DNA in shaping HP1 paralog phase separation and localization. eLife, 2024. 12.

21. Ukmar-Godec, T., et al., Multimodal interactions drive chromatin phase separation and compaction. Proc Natl Acad Sci U S A, 2023. 120(50): p. e2308858120.

22. Roach, R.J., et al., Heterochromatin protein 1alpha interacts with parallel RNA and DNA G-quadruplexes. Nucleic Acids Res, 2020. 48(2): p. 682–693.

23. Wang, L., et al., Histone Modifications Regulate Chromatin Compartmentalization by Contributing to a Phase Separation Mechanism. Mol Cell, 2019. 76(4): p. 646–659 e6.

24. Lou, J., et al., Heterochromatin protein 1 alpha (HP1alpha) undergoes a monomer to dimer transition that opens and compacts live cell genome architecture. Nucleic Acids Res, 2024.

25. Alberti, S., A. Gladfelter, and T. Mittag, Considerations and Challenges in Studying Liquid-Liquid Phase Separation and Biomolecular Condensates. Cell, 2019. 176(3): p. 419–434.

26. Kumar, A. and H. Kono, Heterochromatin protein 1 (HP1): interactions with itself and chromatin components. Biophys Rev, 2020. 12(2): p. 387–400.

27. Walts, J.C. and N.C. Riddle, Phosphorylation as a regulatory mechanism of HP1 protein multifunctionality. Chromosoma, 2025. 134(1).

28. Her, C., et al., Molecular interactions underlying the phase separation of HP1alpha: role of phosphorylation, ligand and nucleic acid binding. Nucleic Acids Res, 2022. 50(22): p. 12702–12722.

29. Narlikar, G.J., Phase-separation in chromatin organization. Journal of Biosciences, 2020. 45(1).

30. Ackermann, B.E. and G.T. Debelouchina, Heterochromatin Protein HP1alpha Gelation Dynamics Revealed by Solid-State NMR Spectroscopy. Angew Chem Int Ed Engl, 2019. 58(19): p. 6300–6305.

31. Chin, S.Y., et al., Nanosecond Molecular Motion in pHP1α Liquid-Liquid Phase Separation Captured by Solid-State NMR. Journal of Physical Chemistry Letters, 2025. 16(4): p. 1150–1156.

32. Chin, S.Y., et al., Investigating Different Dynamic pHP1α States in Their KCl-Mediated Liquid-Liquid Phase Separation (LLPS) Using Solid-State NMR (SSNMR) and Molecular Dynamic (MD) Simulations. Journal of Physical Chemistry B, 2024. 128(42): p. 10451–10459.

33. Chin, S.Y., et al., Role of Trifluoroacetic Acid Anions in pHP1α Liquid-Liquid Phase Separations. Journal of Physical Chemistry B, 2025. 129(31): p. 7964–7971.

34. Kanekura, K., Y. Hayamizu, and M. Kuroda, Order controls disordered droplets: structure-function relationships in C9ORF72-derived poly(PR). Am J Physiol Cell Physiol, 2022. 322(2): p. C197–C204.

35. Solomon, D.A., et al., Altered Phase Separation and Cellular Impact in C9orf72-Linked ALS/FTD. Front Cell Neurosci, 2021. 15: p. 664151.

36. Freibaum, B.D. and J.P. Taylor, The Role of Dipeptide Repeats in C9ORF72-Related ALS-FTD. Front Mol Neurosci, 2017. 10: p. 35.

37. Loveland, A.B., et al., Ribosome inhibition by C9ORF72-ALS/FTD-associated poly-PR and poly-GR proteins revealed by cryo-EM. Nat Commun, 2022. 13(1): p. 2776.

38. Zhang, Y.J., et al., Heterochromatin anomalies and double-stranded RNA accumulation underlie C9orf72 poly(PR) toxicity. Science, 2019. 363(6428).

39. White, M.R., et al., C9orf72 Poly(PR) Dipeptide Repeats Disturb Biomolecular Phase Separation and Disrupt Nucleolar Function. Mol Cell, 2019. 74(4): p. 713–728 e6.

40. Shi, C.W., et al., BSH-CP based 3D solid-state NMR experiments for protein resonance assignment. Journal of Biomolecular Nmr, 2014. 59(1): p. 15–22.

41. Hoffmann, J., et al., Protein resonance assignment by BSH-CP-based 3D solid-state NMR experiments: A practical guide. Magnetic Resonance in Chemistry, 2020. 58(5): p. 445–465.

42. Nowacka, A., et al., Signal intensities in H-C CP and INEPT MAS NMR of liquid crystals. Journal of Magnetic Resonance, 2013. 230: p. 165–175.

43. Reif, B., et al., Solid-state NMR spectroscopy. Nature Reviews Methods Primers, 2021. 1(1).

44. Zeng, T., et al., Characterization of intrinsically disordered regions through scalar coupling-based solid-state NMR experiments. Biophysics Reports, 2025. 11(4).

45. Murthy, A.C. and N.L. Fawzi, The (un)structural biology of biomolecular liquid-liquid phase separation using NMR spectroscopy. J Biol Chem, 2020. 295(8): p. 2375–2384.

46. `Janowska`, M.K. and J. `Baum`, Intermolecular Paramagnetic Relaxation Enhancement (PRE) Studies of Transient Complexes in Intrinsically Disordered Proteins, in Protein Amyloid Aggregation. 2016. p. 45-53.

47. Lenard, A.J., F.A.A. Mulder, and T. Madl, Solvent paramagnetic relaxation enhancement as a versatile method for studying structure and dynamics of biomolecular systems. Progress in Nuclear Magnetic Resonance Spectroscopy, 2022. 132-133: p. 113-139.

48. Shimojo, H., et al., Extended string-like binding of the phosphorylated HP1alpha N-terminal tail to the lysine 9-methylated histone H3 tail. Sci Rep, 2016. 6: p. 22527.

49. Babu, M., et al., Proline/arginine dipeptide repeat polymers derail protein folding in amyotrophic lateral sclerosis. Nat Commun, 2021. 12(1): p. 3396.

50. Wake, N., et al., Expanding the molecular grammar of polar residues and arginine in FUS phase separation. Nature Chemical Biology, 2025. 21(7): p. 1076–1088.

51. Murthy, A.C., et al., Molecular interactions underlying liquid−liquid phase separation of the FUS low-complexity domain. Nature Structural & Molecular Biology, 2019. 26(7): p. 637–648.

52. Usluer, S., et al., p53 Transactivation Domain Mediates Binding and Phase Separation with Poly-PR/GR. Int J Mol Sci, 2021. 22(21).

