## Supplementary material for "Cooperative IDR-folded domain interplay governs HP1α phase separation and underlies biphasic regulation by pathogenic poly-PR": supplymental information

**Fig. S1 |**
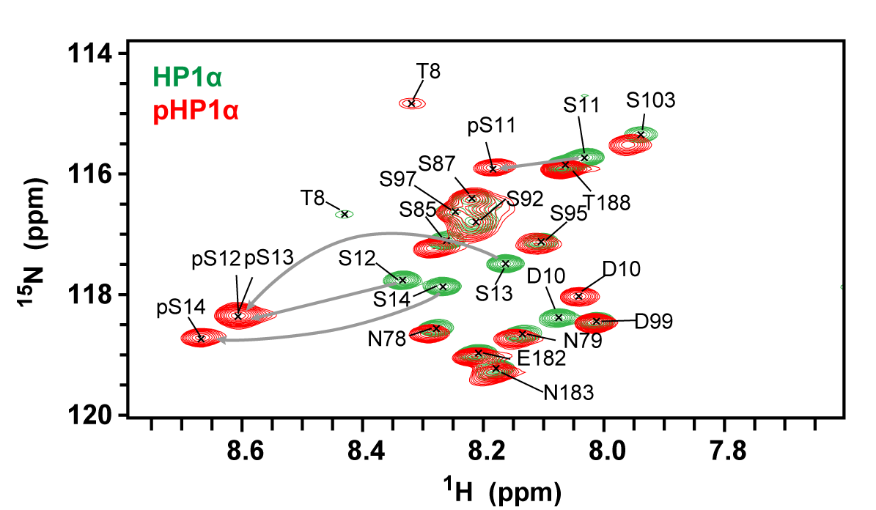
 **Complete phosphorylation of the four NTE serine sites.**

Solution-state ¹H-¹⁵N HSQC spectra of HP1α before and after phosphorylation. The complete chemical shift changes of the S11-S14 resonances corresponding to the NTE phosphorylation sites demonstrate that full phosphorylation has been achieved.

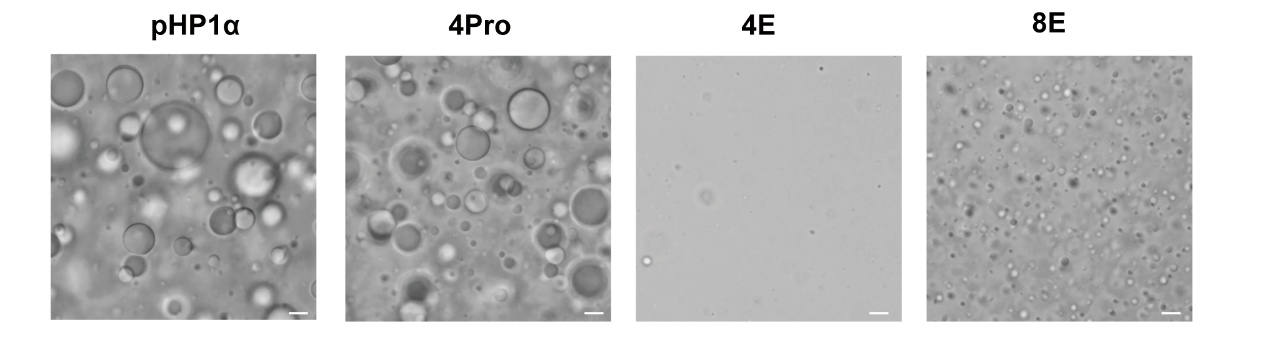
**Fig. S2 | In vitro phase separation of NTE mutants.**

In vitro phase separation microscopy of NTE mutants (4Pro, 4E, and 8E). 4Pro forms droplets comparable to pHP1α, whereas 4E and 8E show minimal or no LLPS. Scale bar: 1 µm

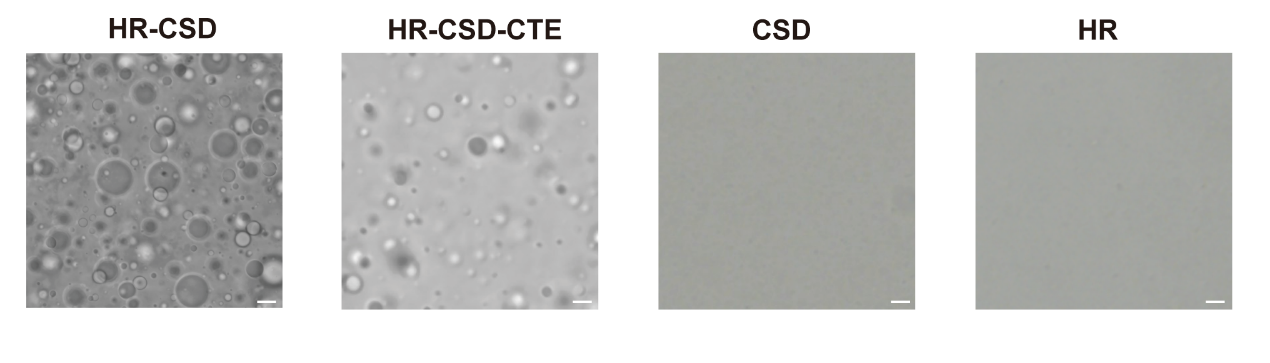
**Fig. S3 | In vitro phase separation of HR-CSD constructs.**

Bright-field microscopy of HR-CSD, HR-CSD-CTE, isolated CSD, and isolated HR. HR-CSD readily undergoes LLPS; HR-CSD-CTE shows reduced LLPS propensity; isolated CSD and HR remain homogeneous without droplet formation. Scale bar: 1 µm.

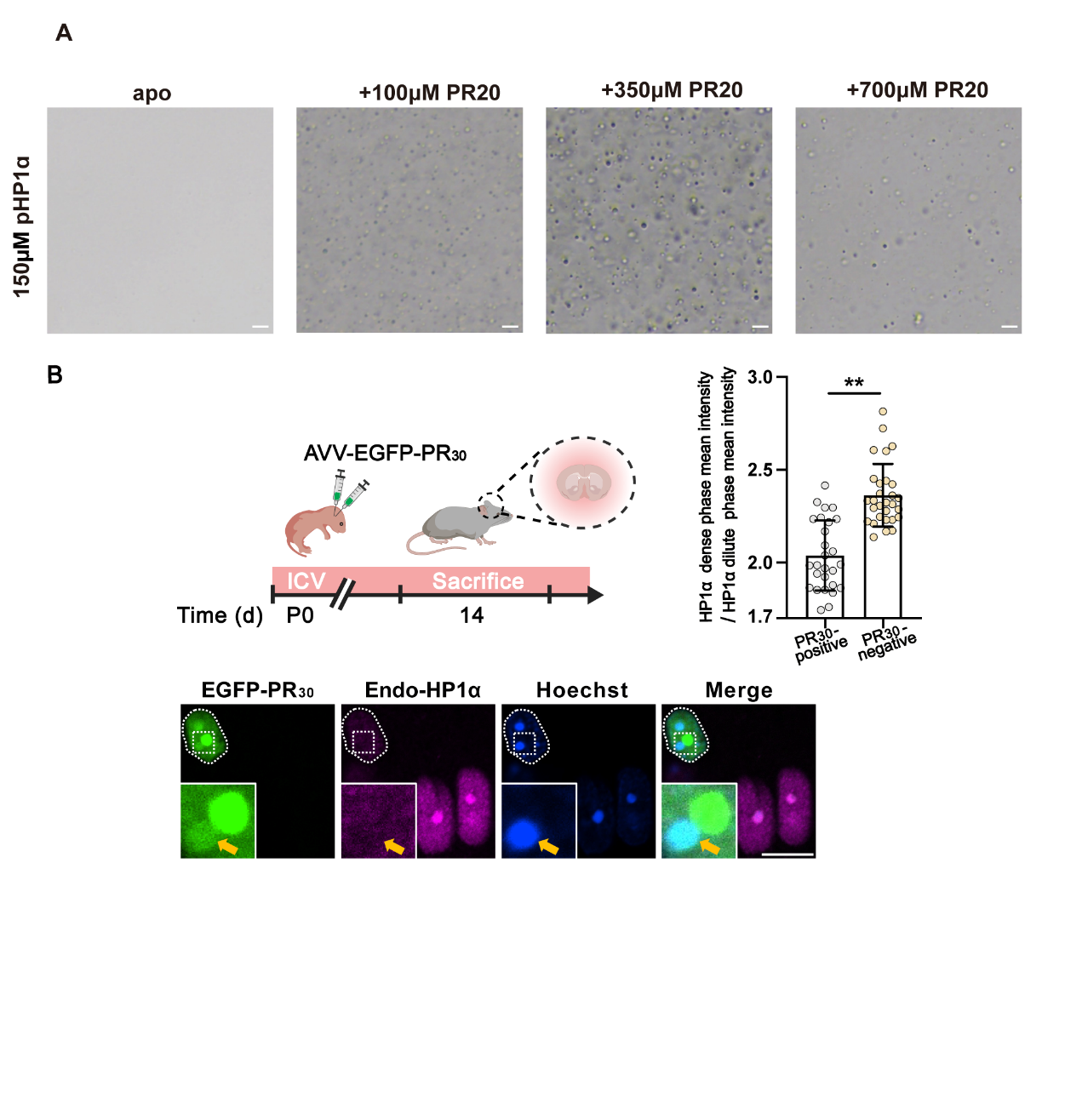
**Fig. S4 | Biphasic regulation of pHP1α LLPS by PR20 in vitro and condensate dissolution under excess poly-PR in vivo.**

**(A)** In vitro phase separation microscopy of 150 µM pHP1α with increasing PR20 concentrations (0, 100, 350, and 700 µM). Low PR20 promotes droplet formation, whereas high PR20 suppresses condensation and reduces droplet size. Scale bar: 1 µm.

**(B)** Brain sections from 14-day-old mice received neonatal injection with AAV-EGFP-PR30. Immunofluorescence shows that PR30-expressing cortical neurons (EGFP-positive) exhibit diffuse endogenous HP1α, whereas neighboring non-expressing cells retain punctate HP1α condensates.

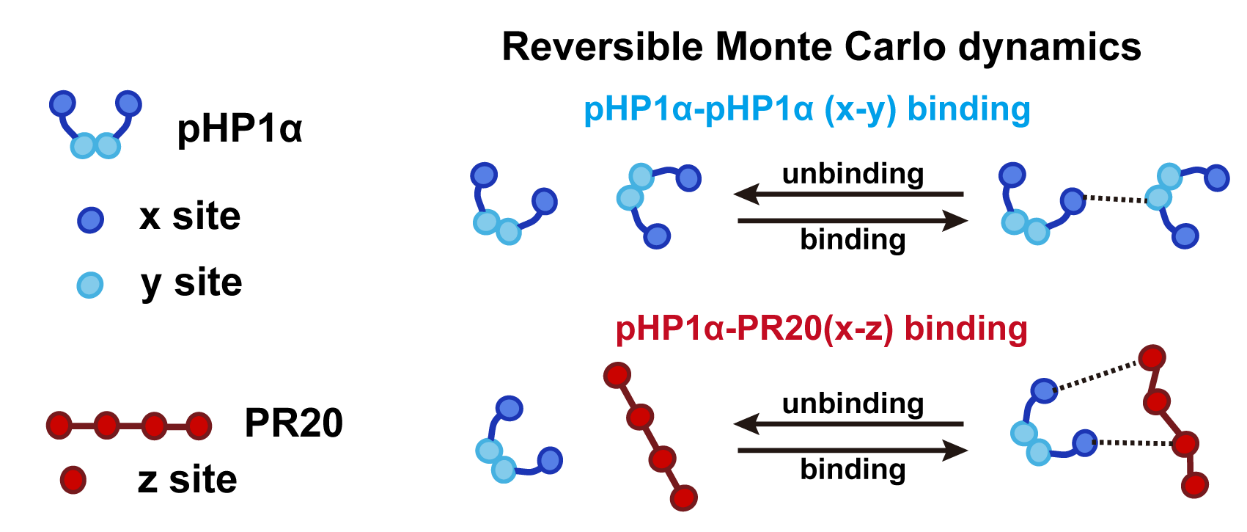

**Fig. S5 | Schematic of the multivalent binding and saturation Monte Carlo (MBS-MC) model.**

pHP1α is represented as a node carrying x and y sites, and PR20 as a node carrying four equivalent z sites. Only two bond types are allowed: x-y bonds, representing basal pHP1α self-association with a weak binding weight, and x-z bonds, representing PR20-mediated crosslinking with a stronger binding weight. Each site participates in at most one bond, imposing strict valency constraints.

**Methods**

**Plasmid Construction**

All plasmids used in this study for non-phosphorylated proteins were cloned into the laboratory-customized pRSFDuet-1 vector, with an N-terminal fusion of a 7×His tag and a TEV protease cleavage site. The plasmids were transformed into BL21 competent cells for protein expression. For phosphorylated proteins, the 6×His-MBP-fused CK2α and CK2β subunit sequences were cloned into the pETDuet-1 vector and co-transformed with the pRSFDuet-1 vector into BL21, as described previously [1-3].

For construction of EGFP-HP1α (WT, 4Pro, 4E and 8E), the coding sequences (HP1α, 4Pro, 4E and 8E) were digested and then cloned into the pEGFP-C1 empty vector through EcoRI/SalI sites, respectively. pmCherry-C1-HP1α was cloned by inserting the coding region of pEGFP-C1-HP1α into a pmCherry-C1 vector plasmid through EcoR I/ Sal I sites.

**Protein Expression and Purification**

For natural-abundance proteins, a single colony from an LB plate was inoculated into 100 mL of LB medium containing the appropriate antibiotic and grown overnight at 37 °C. This culture was then used to inoculate 400 mL of fresh LB medium. When the optical density at 600 nm (OD₆₀₀) reached 0.6-0.8, protein expression was induced by adding 0.5 mM isopropyl β-d-1-thiogalactopyranoside (IPTG), followed by expression for 24 h at 16 °C.

For isotopically labeled proteins, a single colony from an LB plate was first grown in 5 mL of LB medium for 6 h. This starter culture was then transferred into 100 mL of LeMaster & Richards Minimal Medium (LR medium) containing the appropriate antibiotic and grown overnight. The entire 100 mL culture was subsequently used to inoculate 700 mL of fresh LR medium. When the OD₆₀₀ reached approximately 0.8, expression was induced with 0.5 mM IPTG for 24 h at 16 °C. Isotope labeling was achieved by replacing the corresponding components in the LR medium: deuterium labeling used D₂O, ¹⁵N labeling used ¹⁵NH₄Cl as the nitrogen source, and ¹³C labeling used d-[¹³C₆]glucose as the carbon source.

After induction, the cells were harvested by centrifugation. The bacterial pellet was resuspended in 30 mL of binding buffer (20 mM Tris, 300 mM NaCl, 2 mM DTT, pH 7.2) and lysed by high-pressure homogenization. Cell debris was removed by high-speed centrifugation at 19,000 × g. The supernatant was loaded onto a nickel-nitrilotriacetic acid (Ni^2+^-NTA) column (GE Healthcare) pre-equilibrated with binding buffer. The bound protein was eluted with elution buffer (20 mM Tris, 300 mM NaCl, 2 mM DTT, 500 mM imidazole, pH 7.2). The eluate was then subjected to TEV protease cleavage to remove the His-tag, followed by dialysis against 2 L of binding buffer. The dialyzed sample was reloaded onto the Ni-NTA column to remove the TEV protease. For phosphorylated HP1α, this step also removed the His-MBP-tagged CK2α/β subunits used in the phosphorylation reaction. The untagged target protein was collected in the flow-through. Finally, the proteins were concentrated and further purified by size-exclusion chromatography on a Superdex 75 Increase 10/300 GL column (Cytiva) equilibrated with binding buffer. The protein was identified by SDS-PAGE.

H3K9me3, PR20, and PA20 peptides were obtained by solid-phase synthesis (GL Biochem, Shanghai).

**In Vitro HP1α Phosphorylation**

For the isotopically labeled proteins requiring NTE phosphorylation after purification, the protein was phosphorylated using commercial Casein Kinase 2 (CK2) (New England Biolabs). Briefly, 7 μl of CK2 was added to a 1 mL reaction mixture containing 3 mg of the target protein, 33 mM ATP, 50 mM Tris-HCl (pH 7.5), 10 mM MgCl₂, 0.1 mM EDTA, and 2 mM DTT. The reaction was incubated at 30 °C for 12 h. The phosphorylated protein was then separated from the kinase and reaction components by size-exclusion chromatography on a Superdex 75 Increase 10/300 GL column (Cytiva) equilibrated with binding buffer. Phosphorylation efficiency was verified by ¹H-¹⁵N HSQC spectroscopy or SDS-PAGE analysis.

**MTSL Labeling**

As described previously [4], the purified protein sample was first treated with dithiothreitol (DTT) at a final concentration of 500 mM and incubated at 4 °C for 6 h to ensure complete reduction of disulfide bonds. The sample was then buffer-exchanged into MTSL labeling buffer (20 mM Tris-HCl, 150 mM NaCl, pH 7.7) using a PD-10 desalting column (Cytiva). A 100 mM stock solution of (1-oxyl-2,2,5,5-tetramethyl-Δ3-pyrroline-3-methyl) methanethiosulfonate (MTSL) was added to the protein solution at a molar ratio of at least 10:1 (MTSL: protein). The labeling reaction was allowed to proceed in the dark at 4 °C for 12 h. Finally, the labeled protein was dialyzed into the buffer required for subsequent NMR experiments.

**Supernatant Concentration Measurement**

Proteins were concentrated to a starting concentration of 1 mM or 0.95 mM in binding buffer and then dialyzed into phase separation buffer (20 mM Tris-HCl, 75 mM NaCl, 2 mM DTT, pH 7.2) at 4 °C overnight to fully induce phase separation. Following incubation, the samples were centrifuged at 13,000 × g for 10 min at 4 °C. The supernatant was carefully collected, and its protein concentration was measured immediately.

**Phase Diagram Determination**

Prior to experiments, both PR20 and pHP1α proteins were extensively dialyzed against a common buffer containing 20 mM Tris-HCl (pH 7.2), 150 mM NaCl, and 2 mM DTT. This step ensured that the two components were in an identical solution environment, eliminating any buffer‐composition differences that might affect their interaction and phase behavior. After dialysis, the two proteins were mixed at various molar ratios to screen for conditions that induce liquid-liquid phase separation (LLPS). For each mixture, the occurrence and extent of phase separation were evaluated by visual inspection under a bright-field microscope.

For a more quantitative analysis, representative conditions were selected for supernatant concentration measurements. In these titration experiments, the concentration of pHP1α was held constant at a final 150 μM, while PR20 was added in increasing amounts to generate a series of samples. Each 100 μl aliquot of the mixed solution was incubated briefly and then centrifuged at 13,000 × g for 10 min. This centrifugation step pelleted the dense, droplet-rich phase (the condensate), leaving the dilute phase in the supernatant. The protein concentration in the supernatant was then determined using a standard protein assay (e.g., UV absorbance at 280 nm).

**NMR experiments**

All solution NMR experiments were performed on a Bruker AVANCE III 600 MHz spectrometer equipped with a TXI cryogenic probe.

All Solid-state NMR spectra were acquired on a wide-bore Bruker 600 MHz NEO spectrometer using a 1.3 mm HXY probe. The actual sample temperature is determined based on the water peak position.

All NMR spectra were processed with TopSpin 3.6.2 (Bruker) and analyzed with the software package POKY and NMRPipe.

***Sample Preparation***

All solution NMR samples were loaded into standard 5 mm NMR tubes with a sample volume of 400 μl. All solid-state NMR samples were prepared by concentrating the protein to at least 18 mg/mL in 20 mM NaH₂PO₄, 75 mM NaCl, 2 mM DTT, pH 6.5, inducing phase separation at 4 °C, and packing the condensate into a 1.3 mm rotor. For all PRE experiments, DTT was omitted from the buffer to preserve the spin label. Detailed experimental conditions are provided in Supplementary Tables 1, 2, 3.

***Backbone Assignment***

For solution-state NMR, backbone resonance assignments of the target proteins were achieved using uniformly ^13^C-^15^N-labeled samples. A suite of standard triple-resonance experiments was employed, which correlate amide proton and nitrogen frequencies with aliphatic carbon (Cα, Cβ) and carbonyl carbon (CO) chemical shifts. The key experiments included HNCACB and CBCA(CO)NH for connecting sequential residues via Cα/Cβ correlations; HN(CA)CO and HNCO for determining carbonyl carbon assignments; and (H)N(COCA)NH for establishing sequential connectivity through the amide nitrogen. These experiments collectively provide the backbone chemical shift information that underpins the CSP-based interaction mapping and the assignment transfer to solid-state NMR described below.

In solid-state NMR experiments, backbone signal assignment relies on ²H-¹³C-¹⁵N-triple-labeled samples. Signals from the intrinsically disordered region (IDR) are correlated via INEPT transfer pathways, whereas those from the folded domain are transferred using cross-polarization (CP) or SPINE pulse sequences [5]. All experiments were performed under magic‑angle spinning (MAS) conditions to achieve efficient signal detection, thereby yielding site‑specific chemical shifts for backbone atoms.

***NMR Titration Experiments***

For both reference and titration samples, standard ¹H-¹⁵N HSQC spectra were acquired. Combined chemical shift perturbations (CSPs) in the ¹H and ¹⁵N dimensions were calculated using the weighted formula:

$$CSP=\sqrt{\left( \Delta N\times0.1 \right)^{2}+\Delta H^{2}}$$

where ΔH and ΔN are the observed shift changes in ¹H and ¹⁵N, respectively.

Peak intensity ratios were defined as I_b_/I_0_, where I_b_ is the intensity of a residue in the presence of the ligand and I_0_ is its intensity in the reference (ligand-free) spectrum. The associated error (σ) for the intensity ratio I_b_/I_0_ was calculated based on Gaussian error propagation:

$$\sigma=\frac{I_{\text{b}}}{I_{0}}\times\sqrt{\left( \frac{1}{\text{SN}\text{R}_{\text{b}}} \right)^{2}+\left( \frac{1}{\text{SN}\text{R}_{\text{0}}} \right)^{2}}$$

where SNR_b_ and SNR_0_ are the signal-to-noise ratios of the peak in the bound and free states, respectively.

***PRE Experiments***

As described previously [4], for solution NMR PRE measurements, a ¹H-¹⁵N HSQC spectrum was first recorded for the sample containing the MTSL-labeled protein (paramagnetic state). To obtain the diamagnetic reference, 10 µL of 1 M sodium ascorbate (pH 6.5) was added directly into the NMR tube to reduce the spin label. The sample was incubated at room temperature for at least 1 hour to ensure complete reduction before a second ¹H-¹⁵N HSQC spectrum was acquired.

Signal intensity ratios in the PRE experiments were calculated as described in the preceding section.

**Cell culture and plasmid transfection**

N2A (Neuro-2a) cells and HEK 293 cells were cultured at 37 °C in 5% CO2 in Dulbecco’s modified Eagle’s medium (DMEM; Gibco, 11995500) containing 10% fetal bovine serum (FBS; EallBIO, 03.C16001DC) with 1% penicillin-streptomycin (Gibco). Transient transfection was performed using the Hieff TransTM Liposomal Transfection Reagent (Yeasen, 40802ES02) according to the manufacturer’s protocol in Opti-MEM (OMEM; Gibco, 31985070) without serum.

**Immunofluorescence staining**

For immunofluorescence, cells were fixed with 4% paraformaldehyde for 10 min at 4 °C, then washed with PBS three times and permeabilized with 0.1% Triton X-100 in PBS for 6 min. After washing with PBST three times, the cells were incubated with anti-HP1α (Cell Signaling Technology, 2616) at 4 °C overnight, then washed with PBST three times and incubated with secondary antibodies conjugated with Alexa Fluor 594 for 2 h at room temperature, followed by Hoechst 33342 staining for 15 min. After 3 washes with PBS, images were captured by a Nikon Eclipse Ti2 confocal microscope equipped with a 60x oil immersion objective.

Mouse breeding and husbandry were kept in SPF conditions. Anesthetized mice were perfused with chilled PBS and 4% PFA in PBS, and then the brain was carefully dissected. The brains were placed in 4% PFA at 4 °C for 12 h and then stored in 15% sucrose in PBS for 24 h, followed by a replacement with 30% sucrose for another 24 h. PFA-fixed brains were embedded in OCT and cut to a thickness of 20 μm (coronal sections). The sections were washed in PBS three times, then blocked with 1% BSA and permeabilized with 0.3% Triton X-100 for 2 h at room temperature. Anti-HP1α (Cell Signaling Technology, 2616) in PBS containing 1% BSA and 0.3% Triton X-100 was incubated at 4 °C overnight, then the sections were washed with PBS three times and incubated with secondary antibodies conjugated with Alexa Fluor 594 for 2 h at room temperature, followed by Hoechst 33342 staining for 15 min. After 3 washes with PBS, antifade reagent (Solarbio, S2100) was used to mount. The sections were imaged via a Nikon Eclipse Ti2 confocal microscope equipped with a 60x oil-immersion objective.

**Neonatal viral injections**

AAV-PHP.eB-CAG>Kozak-EGFP-PR30 were generated by Cyagen Biosciences Co., Ltd. Intracerebroventricular (ICV) injections of the AAV were performed as previously reported [6, 7]. 2 μL(5E12 GC/mL) of AAV-EGFP-PR30 solution per cerebral ventricle was manually injected into each cerebral ventricle of the pups at postnatal day 0 from a wild-type C57BL/6 mouse.

**Confocal image acquisition and analysis**

Confocal images were monitored with a Nikon Eclipse Ti2 confocal microscope equipped with a 60x oil immersion objective and analyzed using Fiji (NIH). Image processing for 3D reconstruction was performed by Imaris (v. 10.1.0). To obtain the mask of HP1α condensed phase, the image was thresholded via the “Yen” algorithm in Fiji. Then the % HP1α condensed-phase area per nucleus was calculated by dividing the area of the condensed-phase portion by the total nuclear HP1α area.

For mouse brain sections, the integrated intensity of the HP1α dilute phase was obtained by subtracting the integrated intensity of the condensed phase from that of the total nuclear HP1α. The dilute‑phase area was calculated by subtracting the condensed‑phase area from the total nuclear area. The mean intensity of the HP1α dilute phase was then determined by dividing its integrated intensity by its area.

**Multivalent binding and saturation Monte Carlo simulation**

Based on the mechanism suggested by the experiments, we proposed a minimal coarse-grained Monte Carlo model to capture the essential interaction features of pHP1a-PR20 network formation. Our aim was not to represent every microscopic feature of the condensate, but to preserve the key elements required to describe the biphasic response: basal pHP1a-pHP1a association, multivalent PR20-pHP1a binding, and saturation of pHP1a binding sites at high PR20 levels. This modeling strategy allowed us to examine whether PR20-mediated bridging and subsequent site saturation are sufficient to generate the observed biphasic regulation. Accordingly, explicit diffusion, detailed conformational dynamics, and spatially resolved polymer motion were omitted. Phase separation was approximated by the formation of large connected molecular clusters, with network connectivity serving as a proxy for the thermodynamically stable cluster-rich state.

***Coarse-grained protein representation***

In the model, each pHP1a molecule was represented as one particle carrying two distinct binding sites, denoted x and y, whereas each PR20 molecule was represented as one particle carrying four equivalent z sites (Fig. S5). Only two types of bonds were allowed: x-y bonds between pHP1a molecules, representing basal pHP1a self-association, and x-z bonds between pHP1a and PR20, representing PR20-mediated recruitment or crosslinking. No y-z, x-x, y-y, or z-z interactions were permitted.

pHP1α was represented as a mixture of dimeric and tetrameric chains, in accordance with the experimentally observed oligomeric states. The user-defined input specifies the total number of pHP1α monomers and the fraction assigned to each oligomeric state. The number of chains in each state was obtained by dividing the assigned monomer number by its oligomer size and rounding to the nearest integer. A dimeric or tetrameric pHP1α chain carried two or four x sites, respectively, together with the same number of y sites.

***Binding and dissociation***

The simulation was implemented as a well-mixed stochastic binding network. At each Monte Carlo step, the algorithm randomly attempted either bond formation or bond dissociation with equal probability (Fig. S5). For bond formation, one free x site was selected from the pool of unoccupied pHP1a x sites, and one target site was selected from the combined pool of free y sites and free PR20 z sites. If the selected target was a y site, an x-y bond was proposed; if the selected target was a z site, an x-z bond was proposed. Bond formation and dissociation were accepted according to Metropolis-type probabilities determined by the corresponding binding weights, allowing the simulation to sample reversible network assembly rather than irreversible aggregation.

***PR20 occupancy penalty***

Different interaction weights were assigned to the two bond types. The x-y interaction was assigned a weak basal binding probability, reflecting intrinsic pHP1a self-association. The x-z interaction was assigned a stronger PR20-dependent binding weight, reflecting the ability of PR20 to bridge pHP1a molecules. Because each PR20 molecule contains four z sites, a single PR20 chain can connect multiple pHP1a molecules and thereby increase network connectivity. To account for saturation or steric limitation on a PR20 molecule, the effective x-z binding weight was reduced as more z sites on the same PR20 molecule became occupied. This allows the model to capture both multivalent bridging at intermediate PR20 occupancy and loss of productive bridging when PR20 becomes excessive.

***Cluster analysis***

The molecular interaction network was represented as a graph: where pHP1a and PR20 molecules were nodes and accepted x-y or x-z bonds were edges. Condensation was quantified from the connected components of this graph. The largest connected component was used as the main proxy for the cluster-rich state associated with phase separation, and additional network descriptors included the fraction of pHP1a molecules incorporated into the largest cluster, total cluster number, largest cluster size, total occupied x-site fraction, the relative fractions of x-y and x-z bonds, and the fraction of PR20 molecules that bridged two or more pHP1a molecules.

***Parameter scans and data analysis***

Simulations were initialized with specified pHP1α and PR20 molecule numbers, and a fixed system volume was used to convert molecule numbers into normalized concentrations for plotting and comparison. Independent simulations with different random seeds were performed for each parameter combination; replicates were grouped by complete parameter combination and summarized as mean ± sample standard deviation. Parallel processing was used only to run independent trajectories and did not alter the update rules.

***Model interpretation***

Because the model lacks explicit spatial coordinates and solvent, largest-cluster measurements were interpreted as indicators of network formation and dissolution rather than as direct measurements of a physical droplet boundary.

| Supplementary Table 1. ssNMR experimental detail parameters | | | | | | |
| --- | --- | --- | --- | --- | --- | --- |
| Experiment | Real Temperature | MAS | Pulse power | Acquisition time (F1) | Acquisition time (F2) | Acquisition time (F3) |
| pHP1α INEPT ^1^H-^15^N HSQC (pH 7.5) | 298K | 15K Hz | H: 156.25 kHz  N: 55.56 kHz | 39.74ms | 43.01ms | - |
| pHP1α CP ^1^H-^15^N HETCOR | 278K | 55K Hz | H: 161.3 kHz  N: 59.5 kHz | 26.31ms | 14.95ms | - |
| HP1α INEPT HNCO | 298K | 59K Hz | H: 172.41 kHz  N: 55.56 kHz  C: 65.79 kHz | 29.82ms | 29.36ms | 39.98ms |
| pHP1α CP CONH | 278K | 55K Hz | H: 161.29 kHz  N: 59.10 kHz  C: 65.79 kHz | 6.626ms | 9.763ms | 14.95ms |
| pHP1α CP CANH | 278K | 55K Hz | H: 161.29 kHz  N: 59.10 kHz  C: 65.79 kHz | 3.823ms | 9.763ms | 14.95ms |

| Supplementary Table 2. ssNMR experimental detail parameters | | | | | | |
| --- | --- | --- | --- | --- | --- | --- |
| Experiment | CP transfer H→N | CP transfer N→H | CP transfer H→Cα | CP transfer H→Co | SPINE transfer Cα→N | SPINE transfer Co→N |
| pHP1α CP ^1^H-^15^N HETCOR | H: 112.41K Hz  N: 43.79K Hz Ramp: 80- 100 % ramp on H  Contact time: 500us | H: 109.06K Hz  N: 43.79K Hz Ramp: 100- 80 % ramp on H Contact time: 500us | - |  | - | - |
| pHP1α CP CONH | - | H: 109.06K Hz  N: 43.50K Hz Ramp: 100- 80 % ramp on H  Contact time: 500us | - | H: 98.30 K Hz  Co: 40.26K Hz Ramp: 90- 100 % ramp on H Contact time: 2500us | - | N: 6.818us@7.859W  Co: 6.818us@3.953W |
| pHP1α CP CANH | - | H: 109.06K Hz  N: 43.50K Hz Ramp: 100- 80 % ramp on H  Contact time: 500us | H: 102.02K Hz  Cα: 40.25K Hz Ramp: 90- 100 % ramp on H  Contact time: 3500us | - | N: 6.818us@7.859W  Cα: 6.818us@3.953W | - |

| Supplementary Table 3. solution NMR experimental conditions | | | |
| --- | --- | --- | --- |
| Experiment | Temperature | Concentration | buffer |
| pHP1α ^1^H-^15^N HSQC  HP1α ^1^H-^15^N HSQC | 308K | ^15^N pHP1α: 100 μM  ^15^N HP1α: 100 μM | 20 mM NaH₂PO₄, 150 mM NaCl, 2 mM DTT, pH 6.5, with 95% H₂O/5% D₂O |
| HR-CSD-CTE ^1^H-^15^N HSQC  HR-CSD ^1^H-^15^N HSQC | 308K | HR-CSD-CTE: 550 μM  HR-CSD: 500 μM | 20 mM NaH₂PO₄, 150 mM NaCl, 2 mM DTT, pH 6.5, with 95% H₂O/5% D₂O |
| ^14^N HR + ^15^N CSD-CTE ^1^H-^15^N HSQC  ^14^N HR + ^15^N CSD ^1^H-^15^N HSQC | 308K | ^14^N HR: 400 μM  ^15^N CSD-CTE: 100 μM  ^15^N CSD: 100 μM | 20 mM NaH₂PO₄, 150 mM NaCl, 2 mM DTT, pH 6.5, with 95% H₂O/5% D₂O |
| ^14^N CSD + ^15^N CSD ^1^H-^15^N HSQC  ^14^N CSD-CTE + ^15^N CSD-CTE ^1^H-^15^N HSQC | 308K | ^14^N CSD-CTE: 400 μM  ^14^N CSD: 400 μM  ^15^N CSD-CTE: 100 μM  ^15^N CSD: 100 μM | 20 mM NaH₂PO₄, 150 mM NaCl, 2 mM DTT, pH 6.5, with 95% H₂O/5% D₂O |
| ^14^N PR20 + ^15^N pHP1α ^1^H-^15^N HSQC | 298K | ^14^N PR20: 25 μM  ^15^N pHP1α: 50 μM | 20 mM NaH₂PO₄, 150 mM NaCl, 2 mM DTT, pH 6.5, with 95% H₂O/5% D₂O |
| ^14^N PR20 + ^15^N pHP1α ^1^H-^15^N HSQC | 308k | ^14^N PR20: 25 μM  ^15^N pHP1α: 50 μM | 20 mM NaH₂PO₄, 150 mM NaCl, 2 mM DTT, pH 6.5, with 95% H₂O/5% D₂O |
| ^15^N A9C-MTSL NTE-CD ^1^H-^15^N HSQC  ^15^N A9C-MTSL pNTE-CD ^1^H-^15^N HSQC  ^15^N A9C-MTSL 4Pro-NTE-CD ^1^H-^15^N HSQC  ^15^N A9C-MTSL pNTE-CD + ^14^N H3K9me3 ^1^H-^15^N HSQC  ^15^N A9C-MTSL pNTE-CD + ^14^N PR20 ^1^H-^15^N HSQC | 308K | ^15^N A9C-MTSL NTE-CD: 100 μM  ^15^N A9C-MTSL pNTE-CD: 100 μM  ^14^N H3K9me3: 100 μM  ^15^N A9C-MTSL 4Pro-NTE-CD: 100 μM  ^14^N PR20: 100 μM | 20 mM NaH₂PO₄, 150 mM NaCl, pH 6.5, with 95% H₂O/5% D₂O |
| ^14^N HR-CSD-CTE + ^2^H-^15^N S92C-MTSL HR-CSD-CTE ^1^H-^15^N HSQC  ^14^N HR-CSD + ^2^H-^15^N S92C-MTSL HR-CSD ^1^H-^15^N HSQC | 308K | ^14^N HR-CSD: 50 μM  ^2^H-^15^N S92C-MTSL HR-CSD: 50 μM  ^14^N HR-CSD-CTE: 50 μM  ^2^H-^15^N S92C-MTSL HR-CSD-CTE: 50 μM | 20 mM NaH₂PO₄, 150 mM NaCl, pH 6.5, with 95% H₂O/5% D₂O |
| ^15^N S191C-MTSL CSD-CTE ^1^H-^15^N HSQC | 308K | ^15^N S191C-MTSL CSD-CTE: 100μM | 20 mM NaH₂PO₄, 150 mM NaCl, pH 6.5, with 95% H₂O/5% D₂O |

**Reference**

1.	Larson, A.G., et al., *Liquid droplet formation by HP1alpha suggests a role for phase separation in heterochromatin.* Nature, 2017. **547**(7662): p. 236-240.

2.	Furukawa, A., et al., *A dynamic structural unit of phase-separated heterochromatin protein 1α as revealed by integrative structural analyses.* Nucleic Acids Research, 2025. **53**(6).

3.	Ackermann, B.E. and G.T. Debelouchina, *Heterochromatin Protein HP1alpha Gelation Dynamics Revealed by Solid-State NMR Spectroscopy.* Angew Chem Int Ed Engl, 2019. **58**(19): p. 6300-6305.

4.	Janowska, M.K. and J. Baum, *Intermolecular Paramagnetic Relaxation Enhancement (PRE) Studies of Transient Complexes in Intrinsically Disordered Proteins*, in *Protein Amyloid Aggregation*. 2016. p. 45-53.

5.	Xiao, H., et al., *Robust Heteronuclear Correlations for Sub-milligram Protein in Ultrafast Magic-Angle Spinning Solid-State NMR.* Journal of the American Chemical Society, 2025. **147**(8): p. 6384-6389.

6.	Maor-Nof, M., et al., *p53 is a central regulator driving neurodegeneration caused by poly(PR).* Cell, 2021. **184**(3): p. 689-+.

7.	Zhang, Y.J., et al., *Heterochromatin anomalies and double-stranded RNA accumulation underlie C9orf72 poly(PR) toxicity.* Science, 2019. **363**(6428).
